# Adaptive Regularized Tri-Factor Non-Negative Matrix Factorization for Cell Type Deconvolution

**DOI:** 10.1101/2023.12.07.570631

**Authors:** Tianyi Liu, Chuwen Liu, Quefeng Li, Xiaojing Zheng, Fei Zou

## Abstract

Accurate deconvolution of cell types from bulk gene expression is crucial for understanding cellular compositions and uncovering cell-type specific differential expression and physiological states of diseased tissues. Existing deconvolution methods have limitations, such as requiring complete cellular gene expression signatures or neglecting partial biological information. Moreover, these methods often overlook varying cell-type mRNA amounts, leading to biased proportion estimates. Additionally, they do not effectively utilize valuable reference information from external studies, such as means and ranges of population cell-type proportions. To address these challenges, we introduce an Adaptive Regularized Tri-factor non-negative matrix factorization approach for deconvolution (ARTdeConv). We rigorously establish the numerical convergence of our algorithm. Through benchmark simulations, we demonstrate the superior performance of ARTdeConv compared to state-of-the-art semi-reference-based and reference-free methods. In a real-world application, our method accurately estimates cell proportions, as evidenced by the nearly perfect Pearson’s correlation between ARTdeConv estimates and flow cytometry measurements in a dataset from a trivalent influenza vaccine study. Moreover, our analysis of ARTdeConv estimates in COVID-19 patients reveals patterns consistent with important immunological phenomena observed in other studies. The proposed method, ARTdeConv, is implemented as an R package and can be accessed on GitHub for researchers and practitioners.

## 1. Introduction

Heterogeneity in cell-type proportions exists across biological samples, and neglecting this heterogeneity in bulk gene expression can introduce bias into subsequent analyses, such as differential expression analyses. Conversely, acknowledging and accounting for this heterogeneity has shown clear advantages, yielding more accurate survival time predictions and tumor type classifications (Elloumi et al., 2011; Avila Cobos et al., 2020).

Laboratory techniques such as flow cytometry or immunohistochemistry are available for physically sorting cells into cell types and quantifying their abundances. However, these methods are often limited by the availability of cell samples, the specificity of antibodies for separating cells, and the substantial labor and time investments required (Newman et al., 2015; Spitzer and Nolan, 2016). Cell type deconvolution (simply referred to as “deconvolution” in this work), a computational process aimed at digitally separating heterogeneous mixture signals into their constituent components, has been critical in expediting the estimation of cell-type proportions from bulk gene expression data, such as RNA sequencing (RNA-seq). In recent years, several deconvolution methods have emerged, with extensive applications in the field of computational biology. These methods can generally be categorized into two groups: reference-based and reference-free, depending on whether they require individual cell-type gene expression signatures, referred to as a signature matrix, as prior knowledge (Li and Wu, 2019; Avila Cobos et al., 2020; Jin and Liu, 2021).

Gene signatures can be derived from either single-cell RNA-seq (scRNA-seq) data or sorted bulk RNA profiles of individual cell types and used as a reference (Chen et al., 2018). Although gene signatures have been successfully established for some cell types, acquiring those for other cell types might be labor-intensive or even unfeasible (Morris and Kopetz, 2016; Wigerblad et al., 2022). Neglecting to incorporate gene signatures for a prevalent cell type can induce biases in the proportional estimates of other cell types within reference-based deconvolution (Avila Cobos et al., 2020). On the other hand, reference-free techniques enable the unsupervised estimation of cell-type proportions but at the cost of disregarding information embedded in known gene signatures (Zaitsev et al., 2019; Chen et al., 2020). A deconvolution method that can utilize partial reference gene signatures presents a plausible compromise.

Additionally, external information on population cell-type proportions may also aid in estimating cell-type proportions. For example, in blood samples, the proportion means (or medians) and ranges of major cell types are available from complete blood count (CBC) tests. However, deconvolution techniques that effectively incorporate such information are currently lacking in the field.

During data pre-processing, information specific to the quantities of RNA molecules contained within an individual cell is often lost after library normalization. Zaitsev et al. (2019) found that failures to account for mRNA amounts produced biased cell-type proportion estimates. However, many deconvolution methods assume a generative model where the loss of this information cannot be accounted for (Avila Cobos et al., 2020). Therefore, incorporating these quantities into the generative model can be pivotal to deconvolution accuracy.

In recent years, several semi-reference-based methods have been developed to perform deconvolution using only partial references. EPIC addresses the lack of representation of cancer cells in common gene signature matrices for the deconvolution of the tumor microenvironment (TME) (Racle et al., 2017; Racle and Gfeller, 2020). Specifically, EPIC treats all types of cancer cells as a single super cell type in deconvolution, allowing for a detailed characterization of immune cell proportions in the TME while accounting for the fraction of cancer cells. This super-cell-type rationale can also be extended to the deconvolution of other tissue types, such as blood. Moreover, EPIC assigns weights to each gene in the signature matrix based on its variability across cell types; genes with lower expression variance are given higher weights for deconvolution. EPIC also adjusts for differing mRNA amounts between cell types, which must be specified prior to deconvolution. However, this becomes infeasible if the tissue is complex, especially when there is a super cell type with multiple constituent cell types whose reference expression is missing. EPIC further requires that deconvolution be performed using signature genes not specific to the uncharacterized super cell type and estimates the proportions of cell types with reference using an inequality-constrained optimization procedure. Consequently, the estimated proportions of the cell type with a missing reference are biased toward zero (Tai et al., 2021).

Two other tools, quanTIseq and SECRET, have similar functions (i.e., they can accommodate one cell or super cell type with a missing reference), with the former essentially employing the same mathematical model as EPIC (Finotello et al., 2019; Lu et al., 2023). An innovation of quanTIseq over EPIC is its carefully defined set of reference gene signatures for immune cells and the calculation of cell-type mRNA amount scaling factors using housekeeping genes (Finotello et al., 2019), but it does not weigh the genes for deconvolution. These innovations also render quanTIseq unable to use customized signature matrices when needed. On the other hand, SECRET can incorporate customized gene signatures from scRNA-seq experiments and uses an outlier-insensitive *L*_1_ norm on the residuals instead of EPIC’s *L*_2_ norm (Lu et al., 2023). However, it does not truly utilize any reference information about the uncharacterized cell type, making it prone to the same bias toward zero as occurs with EPIC.

BayICE is another semi-reference-based method serving the same purpose as the other methods above, but it adopts a hierarchical Bayes design with stochastic gene signature selection (Tai et al., 2021). Due to its probabilistic nature, BayICE can quantify the uncertainty of cellular abundance estimates and claims to mitigate the biases of EPIC. These benefits come at the expense of increased computational complexity, sensitivity to priors, and difficulty in interpreting the parameters involved. Moreover, it requires bulk samples of purified cells as reference inputs, which, like quanTIseq, precludes the use of other types of reference data, such as customized signature matrices.

In this article, we propose a novel semi-reference-based method called ARTdeConv (short for “Adaptive Regularized Tri-factor non-negative matrix factorization method for deConvolution”). It addresses the three outstanding issues discussed above: utilizing partial reference gene signatures, incorporating external information on cell-type proportions, and accounting for cell-type mRNA amounts. Compared to EPIC, ARTdeConv does not require the specificity of the gene signatures to cell types that are characterized. It also uses external proportion information to correct for EPIC’s observed biases toward zero. Moreover, compared to EPIC and quanTIseq, ARTdeConv learns the relative cell-type mRNA abundances automatically without the need to use housekeeping genes or other information while maintaining the straightforward interpretation of the resulting estimates. It can also incorporate scRNA-seq data as SECRET does and uses a different optimization procedure. Our further contributions include deriving and implementing a multiplicative update (MU) algorithm for solving ARTdeConv, proving the algorithm’s convergence, and demonstrating its merits through simulations and real data analysis. A schematic representation of ARTdeConv’s workflow is shown in Figure 1. An R package implementing ARTdeConv is also available on GitHub.

**Figure 1:**
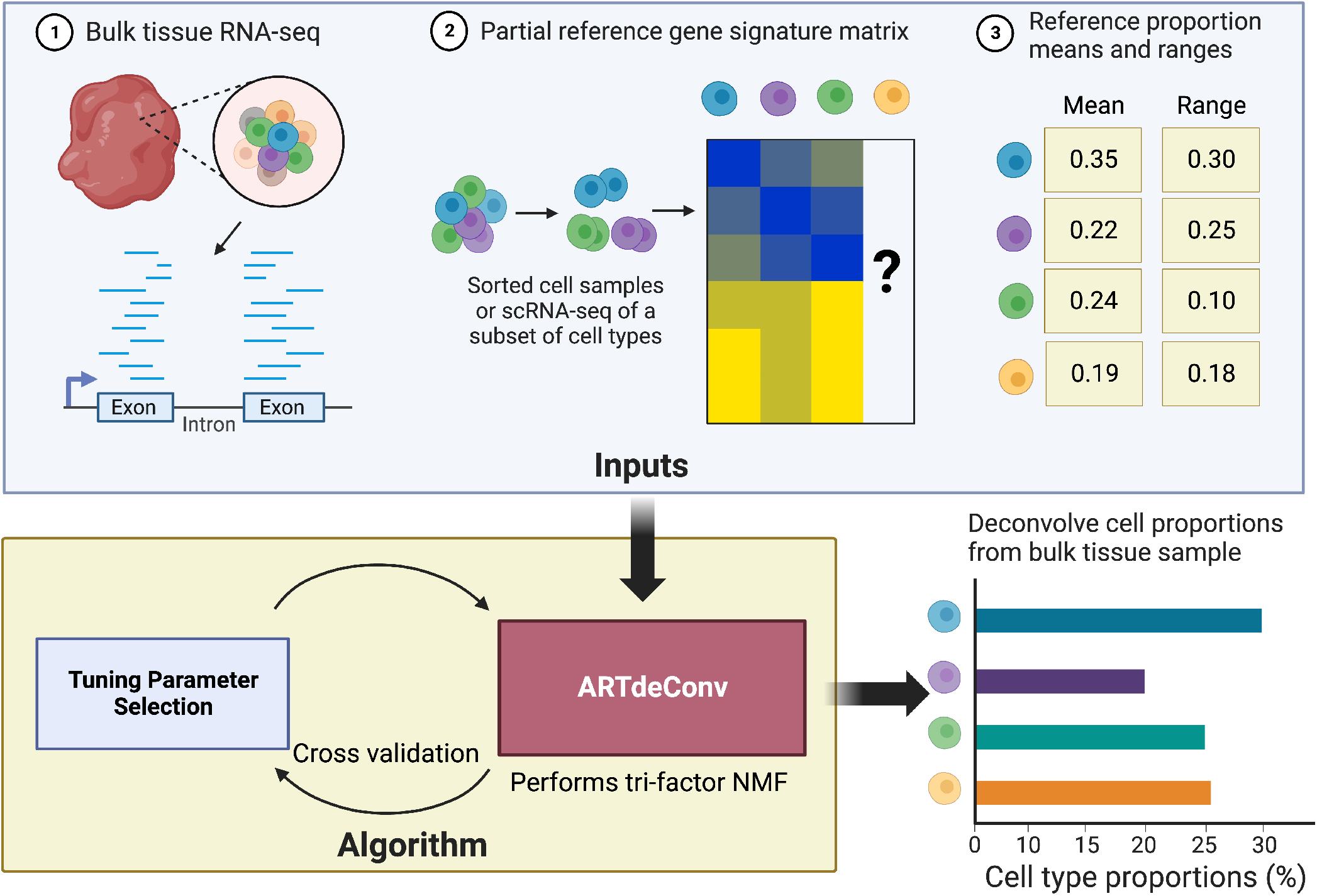
A schema of the ARTdeConv workflow. Specifically, ARTdeConv takes in three parts of information: the gene expression data of the bulk tissue to be deconvolved, the reference gene expression data with one cell type or super cell type uncharacterized, and the reference means/medians and ranges of the proportions of all cell types in deconvolution. It then passes these inputs to a cross validation algorithm to select the optimal hyperparameters. Lastly, these hyperparameters are passed along the rest of the inputs for a deconvolution run that estimate the proportions of each cell type or super cell type involved.

## 2. Method

### 2.1. Notation

Let ℝ_+_ and ℝ_++_ denote the set of non-negative and positive real numbers respectively. *m, n, K* are positive integers used to denote the number of genes, samples, and cell types with *K* ≤ min(*m, n*) in deconvolution. Let a positive integer *K*_0_ denote the number of cell types for which we have reference gene expression available. Unless otherwise mentioned, we set *K* = *K*_0_ + 1. Let 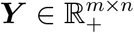 the matrix of bulk gene expression and 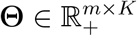 be the full gene signature matrix. Let 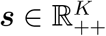 be a vector of relative cell-type mRNA amounts, and 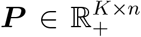 the proportion matrix. Let ***ϵ*** ∈ ℝ^*m×n*^ be the random error matrix. Unless otherwise specified, we denote ***θ***_*k*_ as the *k*-th column of **Θ**. Let ***A*** be a genereic *m* × *n* matrix. We use ***A***^⊤^ to denote the transpose of a matrix. If ***A*** is further a square matrix (*m* = *n*), we denote its trace 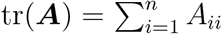. For any *m* × *n* matrix ***A***, its Frobenius norm is denoted as 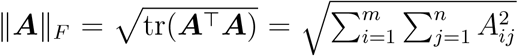. For two matrices ***A, B*** of the same dimensions, ***A*** ⊙ ***B*** denotes the element-wise product of ***A*** and ***B*** and 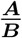 denotes their element-wise quotient.

### 2.2. Model And Problem Setup

We first propose the following tri-factor generative model

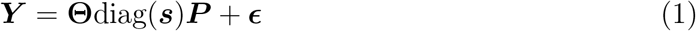

for ARTdeConv, which extends the canonical model ***Y***
= **Θ*P*** + ***ϵ*** (Avila Cobos et al., 2020). The difference between them is diag(***s***), a diagonal matrix of cell-type mRNA amounts. In practice, normalization (e.g. by library sizes, etc.) is frequently employed to alleviate between-sample technical artifacts in sequencing data (Hafemeister and Satija, 2019; Lytal et al., 2020). During this process, information regarding the quantities of RNA molecules packed within different kinds of cells sometimes gets lost. Failures to recover these quantities lead to documented biases in estimating cell-type proportions (Racle et al., 2017; Zaitsev et al., 2019). Should this occurs, diag(***s***) can account for the RNA molecule quantities.Otherwise, diag(***s***) would be close to the identity matrix ***I***_*K*_ (an option to fix diag(***s***) = ***I***_*K*_ is given by ARTdeConv). The values in diag(***s***) should be interpreted in a relative sense. For example, *s*_2_*/s*_1_ is the ratio between the amounts of mRNA molecules packed by the second and the first cell type. We do not require any prior specification of diag(***s***) and notice that (1) is similar to the model of MuSiC, a reference-based method where each sample is accorded an additional scaling factor *c*_*i*_ to account for between-sample measurement variation in bulk gene expression (Wang et al., 2019). Moreover, this setup differs from that of EPIC, which requires ***s*** be manually specified (Racle et al., 2017).

For performing deconvolution based on (1), ARTdeConv assumes prior knowledge of a partial signature matrix 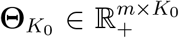 as well as ***Y***. Then, an objective function is defined:

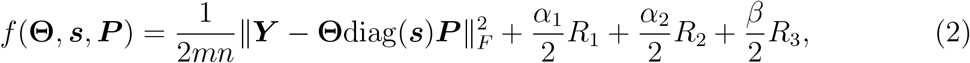

where *R*_1_, *R*_2_, and *R*_3_ are regularizers to be explained later and *α*_1_, *α*_2_, and *β* are their tuning parameters. We discuss the selection of optimal tuning parameters in Section 2.4. **Θ, *s***, and ***P*** are then estimated via

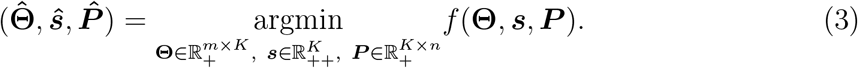

The main objective here is to obtain an estimate of the proportion matrix 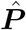. To follow a common practice of deconvolution and simplify the algorithm (Mohammadi et al., 2017), ARTdeConv does not directly constrain each column of ***P*** to sum to one during the estimation process. Instead, it obtains an unconstrained estimate 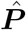 and then re-normalizes its columns to have the unit sum.

Let 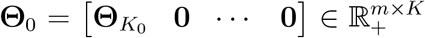 and **Δ** be an *m* × *K* matrix such that Δ_*jk*_ = *I*(*k* ≤ *K*_0_) for 1 ≤ *j* ≤ *m*. The squared Frobenius distance between the estimated **Θ** and **Θ**_0_ is penalized through 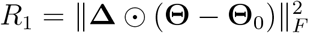. Though we present a special case where 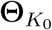 occupies the first *K*_0_ columns of **Θ**, by re-defining **Δ** correspondingly, 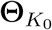 can occupy any *K*_0_ columns, covering all structures of prior knowledge on the signature matrix. Same as other well established semi-reference-based methods (i.e., EPIC, quanTIseq, etc.), we set *K* = *K*_0_ + 1. We also recommend that the unrepresented cell types in 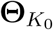 be grouped into a single artificial “cell type”. Albeit that ARTdeConv technically allows *K > K*_0_ + 1, we have found that the resulting estimates from ARTdeConv became less reliable as *K − K*_0_ grew larger in preliminary analysis (results not shown). This makes ARTdeConv suitable for the situations where a major cell type is missing from the reference or that distinguishing the multiple cell types that are missing is not salient. We shall see such examples at work late in Sections 3.2 and 3.3.

On the other hand, *R*_1_ is not a strictly convex function of **Θ**, making the guarantee of the numerical convergence of ARTdeConv’s algorithm difficult (more details on this later). To atone for this, we make 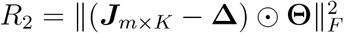, where ***J***_*m×K*_ is an *m* × *K* matrix of 1s. It can be shown that any positive linear combination of *R*_1_ and *R*_2_ is strictly convex with respect to **Θ**. It is of notes that *R*_2_ will force the estimated gene signatures of the uncharacterized close to zero. While this is the case for genes that are specific to the cell types in 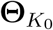, ARTdeConv does not restrict the gene signatures to only those genes. Thus, it is recommended to reduce the penalization effects *R*_2_ by setting its tuning parameter *α*_2_ to a very small value.

Information on the cell-type proportions such as means (or medians) and ranges in a population frequently exists in external data and is accessible online, for example through complete blood counts of leukocytes (Takami et al., 2021). Let *m*_*k*_ denote the reference mean (or median) of cell type *k*’s population proportion and let *r*_*k*_ denote its range. To incorporate it into our deconvolution method, we propose adding another penalty 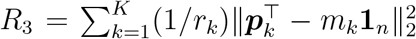, where 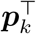 is the *k*-th row of ***P***. In *R*_3_, *m*_*k*_ acts as a pivot, and the deviation from which is penalized for the estimated 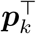. Meanwhile, 1*/r*_*k*_ acts as a weight so the proportion estimates of cell types with wider ranges are less penalized for departing from their respective *m*_*k*_. Letting ***M*** be a *K* × *n* matrix such that ***M*** = [*m*_1_**1**_*n*_ *m*_2_**1**_*n*_ … *m*_*K*_**1**_*n*_]^T^ and ***ρ*** = diag(*r*_1_, *r*_2_, …, *r*_*K*_), we can write *R*_3_ in a matrix form 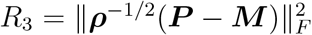.

### 2.3. Multiplicative Update Algorithm

A multiplicative update (MU) algorithm is proposed to solve (3). The MU algorithm was originally designed to solve the canonical bi-factor non-negative matrix factorization (NMF) problem (Lee and Seung, 2000) and was extensively studied (Lin, 2007). Technically, it is similar to the majorization-minimization algorithm of Lange (2016). It can be readily extended to solving a multi-factor regularized NMF problem like the one for ARTdeConv.

Let *t* be a non-negative integer denoting the current number of iteration. The multiplicative update steps are derived by finding the gradients of a set of auxiliary functions for each row of **Θ**^*t*^, each column of ***P*** ^*t*^, and ***s***^*t*^. These update steps are:

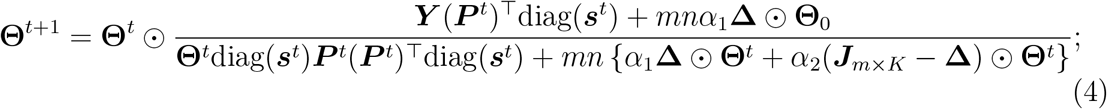

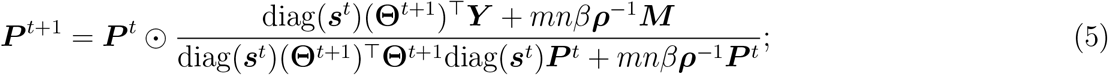

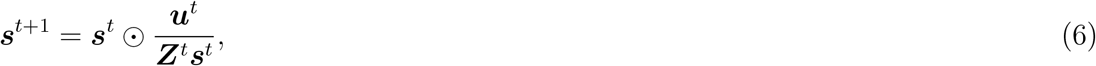

where 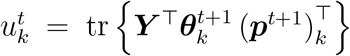 and 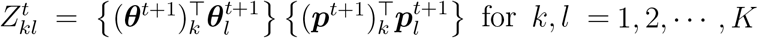.

Mathematical details for finding the auxiliary functions and the update steps can be found in Supplementary Material Section B. The pseudo-code of the MU algorithm is shown in Algorithm 1.

#### Algorithm 1. The MU Algorithm For Solving ARTdeConv

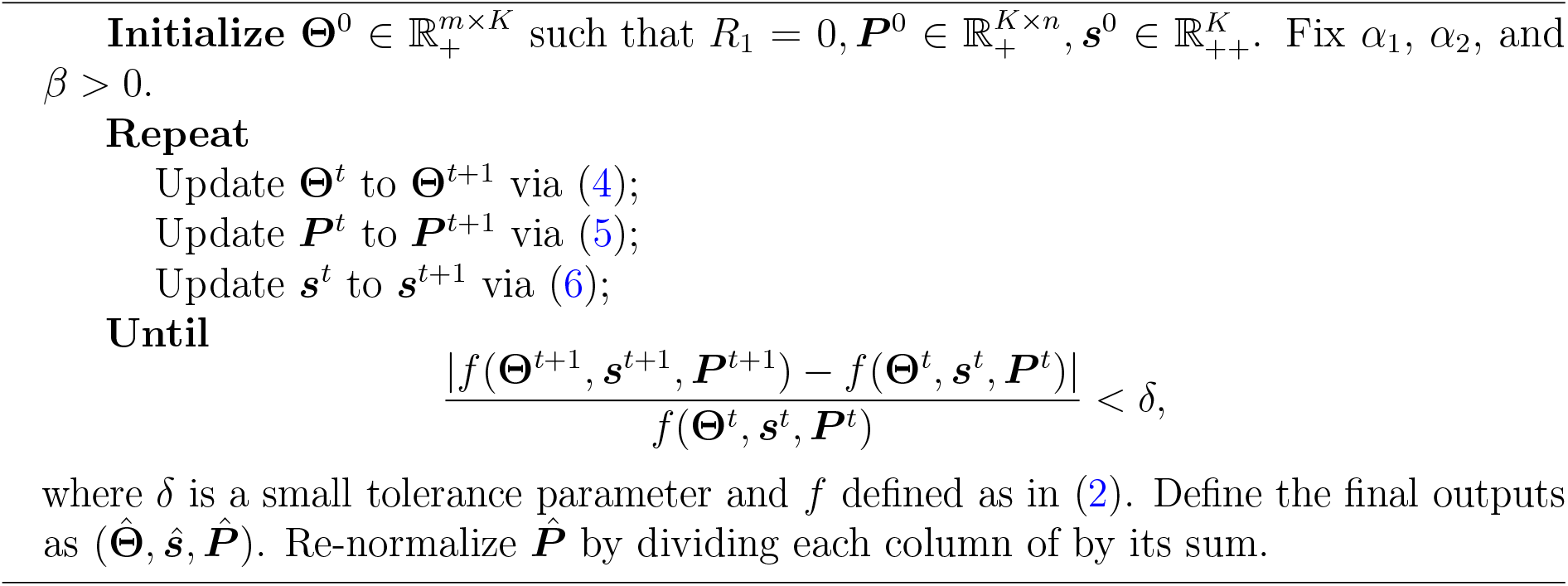

The MU algorithm has two advantages (under the assumptions discussed in Supplementary Material Section A). First, **Θ**^*t*^, ***P*** ^*t*^, and ***s***^*t*^ remain non-negative throughout all iterations if their initial values are non-negative. It is recommended that **Θ**^0^ is set to satisfy *R*_1_ = 0, so any zero in the known partial signature matrix will remain zero. Second, the MU algorithm can be shown to achieve numerical convergence. If we enforce diag(***s***^*t*^) = ***I***_*K*_ and set *α*_1_ = *α*_2_ = *β* = 0, our MU algorithm is reduced to that of Lee and Seung (2000) for the canonical NMF. Due to the non-linearity of the objective function (2), it is not guaranteed that a global minimizer can be produced. Following the precedence of other NMF-based deconvolution software, we recommend restarting the MU algorithm multiple times and choosing the run with the smallest Frobenius norm of the residuals on the estimated bulk expression (Gaujoux and Seoighe, 2010, 2013). We set 10 restarts as the default in our implementation.

We now describe the default initialization procedure of ARTdeConv. First, the first *K*_0_ columns of **Θ**^0^ were used as 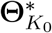 to satisfy *R*_1_ = 0. Each initial value in the last column 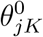 is generated independently by 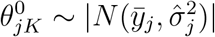, where 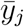 and 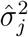 are the mean and variance of the bulk expression values for gene *j* across all *n* samples. This differs from the procedure of SECRET (Lu et al., 2023), which initiates 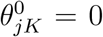. This is because the MU algorithm would produce 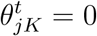 for any *t* = 1, 2, …, *T* following this initiation. Then, for each cell type *k* of sample *i*, 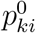 is generated from a *N* (*m*_*k*_, 0.1) distribution. After this, negative entries in ***P*** ^0^ are corrected to 0.01. Lastly, ***s***^0^ is initialized by solving

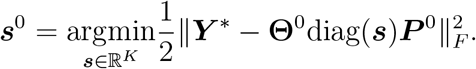

Negative values in ***s***^0^ will be replaced by the mean of positive values.

### 2.4. Tuning Parameter Selection

A grid search using a *B*-fold cross-validation (CV) is designed for selecting tuning parameters. Since *R*_2_ is designed to make the objective function in (2) strictly convex, *α*_2_ is advised to be fixed at a minuscule value. In the implementation of ARTdeConv, *α*_2_ = 10^−12^ is the default value.

To begin, ***Y*** is randomly divided into *B* different folds, each containing 1*/B* columns of ***Y***. Two grids **𝔄**_1_, **𝔅** ⊂ R_++_ containing respective candidates values for *α*_1_ and *β* are declared. Next, for one *α*_1_ ∈ **𝔄**_1_, one *β* ∈ **𝔅** and a fold *b* ∈ {1, 2, …, *B*}, the columns in the fold are held out as the test set 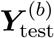. The rest of the columns in ***Y*** are used as the training set 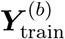, upon which an ARTdeConv solution 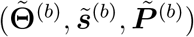 is obtained.

Then, since it is assumed that *n* samples should share the same **Θ** and ***s*** in (1), the estimated proportions 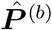 on the test set 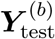 is obtained by

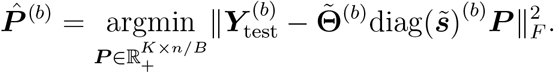

Computationally, this is accomplished by solving a non-negative least square problem using the R package nnls. The CV error given *α*_1_ and *β* is

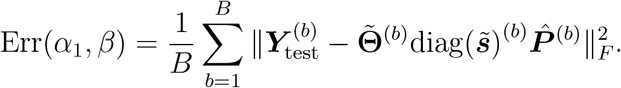

Finally, we select the best tuning parameters 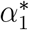 and *β*^*^ that minimize Err(*α*_1_, *β*).

### 2.5. Convergence Analysis

In this section, we prove the numerical convergence of Algorithm 1. The main result depends on two reasonable assumptions in Supplementary Material Section A. Consequently, the following theorem holds for Algorithm 1 for solving the problem in (3).

#### Theorem 1

(Convergence of Algorithm 1 to a Stationary Point). *Under the technical assumptions listed in Supplementary Material Section A, the sequence* 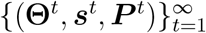 *in Algorithm 1 converges to a stationary point of f* (**Θ, *s, P***).

The proof of Theorem 1 relies on showing that it is a special case of the block successive upper-bound minimization (BSUM) algorithm given in Razaviyayn et al. (2013). Thus, the convergence is guaranteed by related BSUM theories. Doing so involved proving that the sub-level set χ^0^ = {(**Θ, *s, P***) : *f* (**Θ, *s, P***) ≤ *f* (**Θ**^0^, ***s***^0^, ***P*** ^0^)} is compact and the objective function in (3) is coercive, as well as demonstrating the majorization-minimization (MM) properties and strong convexity of the auxiliary functions used to derive the update steps in Algorithm 1. Details on the definitions of coercive functions and MM properties, as well as the proof of Theorem 1 are relegated to Supplementary Material Section C.

## 3. Results

### 3.1. Deconvolution Performance Benchmarks on Pseudo-Bulk Samples

To assess the deconvolution performance of ARTdeConv in comparison to alternative methods, we conducted a benchmarking simulation study by evaluating it against two semi-reference-based methods: EPIC and SECRET, and three reference-free methods: NMF (Gaujoux and Seoighe, 2010), debCAM (Chen et al., 2020), and LINSEED (Zaitsev et al., 2019). We deemed the NMF application here semi-supervised and called it “Semi-NMF” due to the prior knowledge on **Θ**, which was used as the initial values for the basis matrix in NMF. BayICE and quanTIseq were not included due to their required inputs’ incompatibility with the simulation setup.

We generated a set of pseudo-bulk expression matrices using methodologies similar to those outlined in other previous studies (Avila Cobos et al., 2020) by the following formula

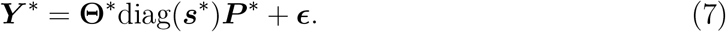

Here, simulated cellular gene expression for *K* = 5 hypothetical cell types, CT1 - CT5, were constructed with assumed prior knowledge on *K*_0_ = 4 of them (CT1 - CT4). We also directly simulated the true signature matrix **Θ**^*^, as opposed to using aggregated purified bulk data or single cell data, for mimicking the widely common practice of using pre-constructed signature matrices in deconvolution applications (Newman et al., 2015; Monaco et al., 2019).

In total, *n* = 200 samples were simulated on ***P*** ^*^. To investigate the effects of the true relative abundance of CT5 in bulk tissue samples on deconvolution results, three classes of ***P*** ^*^ were generated, representing when CT5 was rare, uniform, and extra compared to other cell types in the tissue. Reference medians (*m*_1_, …, *m*_5_) and ranges (*r*_1_, …, *r*_5_) were obtained directly from each row of ***P*** ^*^.

The expression values in the true signature matrix **Θ**^*^ were simulated row-by-row (i.e., gene-by-gene), controlled by a parameter *γ*, which dictated how cell-type-specific each gene was on average. Each simulated gene was more likely to be cell-type-specific when *γ* = 1 compared to when *γ* = 0. The expression of *M* = 2, 000 genes was first created in a matrix 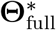. Then, from these genes, a subset of *m* = 1, 000 highly cell-type-specific genes called marker genes were selected for ARTdeConv, Semi-NMF, EPIC, and SECRET. In addition, ***s***^*^ (including 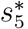) was given to EPIC as for adjusting the mRNA amount per cell type. The method for the selection is described in details in Supplementary Material Section D.2. The expression of those selected genes was stored in **Θ**^*^. The matrix 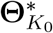 consisting of the first four columns of **Θ**^*^ was used as the partial signature matrix for ARTdeConv, EPIC, and SECRET, and the initial value in the basis matrix for Semi-NMF. Each gene was given the same unit weight in EPIC and SECRET during simulation, for all of the simulated gene expression were derived from the identical generative process. On the other hand, debCAM and LINSEED searched for marker genes differently from 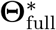 by looking for simplicial vertices of the vector space spanned by normalized bulk gene expression (Zaitsev et al., 2019; Chen et al., 2020).

The matrix of errors ***ϵ*** was first generated for all of the *M* = 2, 000 simulated genes. Beginning with (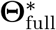, ***s***^*^, ***P*** ^*^), the error-free full bulk expression matrix was calculated as 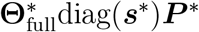. The relative strength of the errors (noises) to the mean expression of genes (signals) was controlled by another parameter *σ*. Two levels of noises with *σ* = 0.1 or 10 (low vs. high) were introduced to evaluate the robustness of the methods to added noises. The final bulk expression matrix of all simulated genes was calculated using 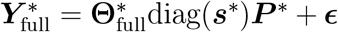. From this, a sub-matrix ***Y*** ^*^ corresponding to the genes in **Θ**^*^ was selected. A detailed description of the probability distributions and their parameters in the generation of **Θ**^*^, ***s***^*^, ***P*** ^*^, and ***ϵ*** can be found in Supplementary Material Section D.1.

With necessary parts generated, 100 simulations were conducted for each combination of *γ, σ*, and CT-5 abundance class, where the deconvolution performance of all benchmarked methods was reported. The tuning grid and algorithm parameters for ARTdeConv are described in Supplementary Material Section D.3. Due to a lack of identifiability, estimated proportions from Semi-NMF, debCAM, and LINSEED were manually matched to different cell types, whose details were relegated to Supplementary Material Section D.4.

Performance metrics for evaluating deconvolution results included the following: (a) 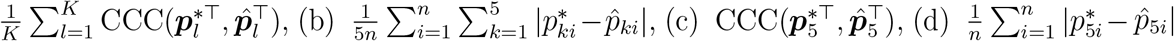, where CCC denotes the Canonical Correlation Coefficient (Lin, 1989; Huang et al., 2023) between two vectors. An advantage of CCC over Pearson’s correlation is that CCC directly measures the agreement between two sets of values by penalizing deviations from the 45-degree line in a scatterplot. Among the four metrics, (a) and (b) delineated the overall deconvolution accuracy, while (c) and (d) described the deconvolution accuracy for CT5, the missing cell type.

Overall, ARTdeConv demonstrated superior performance compared to other semi-reference-based and reference-free methods in accurately recovering cell-type proportions, irrespective of the cell-type specificity of signature genes, the level of additive noise, and the relative abundance of the missing cell type compared to other cell types (Figures 2a and 3a). Notably, ARTdeConv also exhibited robust performance in estimating the proportions of CT5, especially when CT5 was relatively prevalent in the pseudo-bulk samples under high noise conditions, where other methods, except debCAM, showed reduced precision (Figures 2b and 3b).

**Figure 2:**
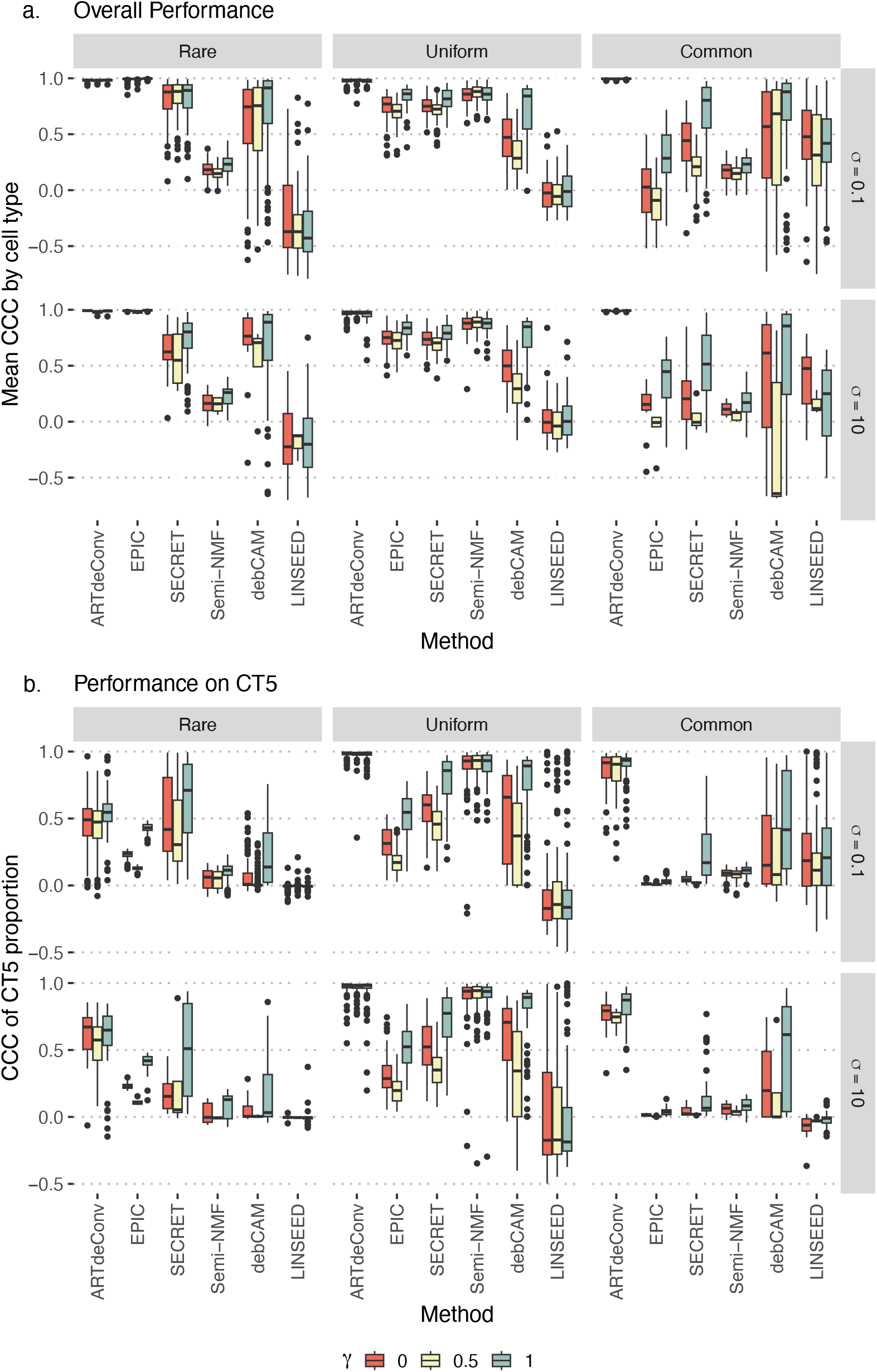
**a**. Mean Concordance Correlation Coefficient (CCC) by cell type between true proportions and estimated cell-type proportions from simulated pseudo-bulks. **b**. CCC between estimated and true proportions of CT5 in the pseudo-bulks for the benchmark simulations. Each column represents a case of the relative abundance of CT5 against other cell types in the simulated pseudo-bulks. Each row represents the level of noises controlled by *σ*. Colors represent different levels of cell-type specificity of genes in **Θ**^*^ regulated by different *γ*.

**Figure 3:**
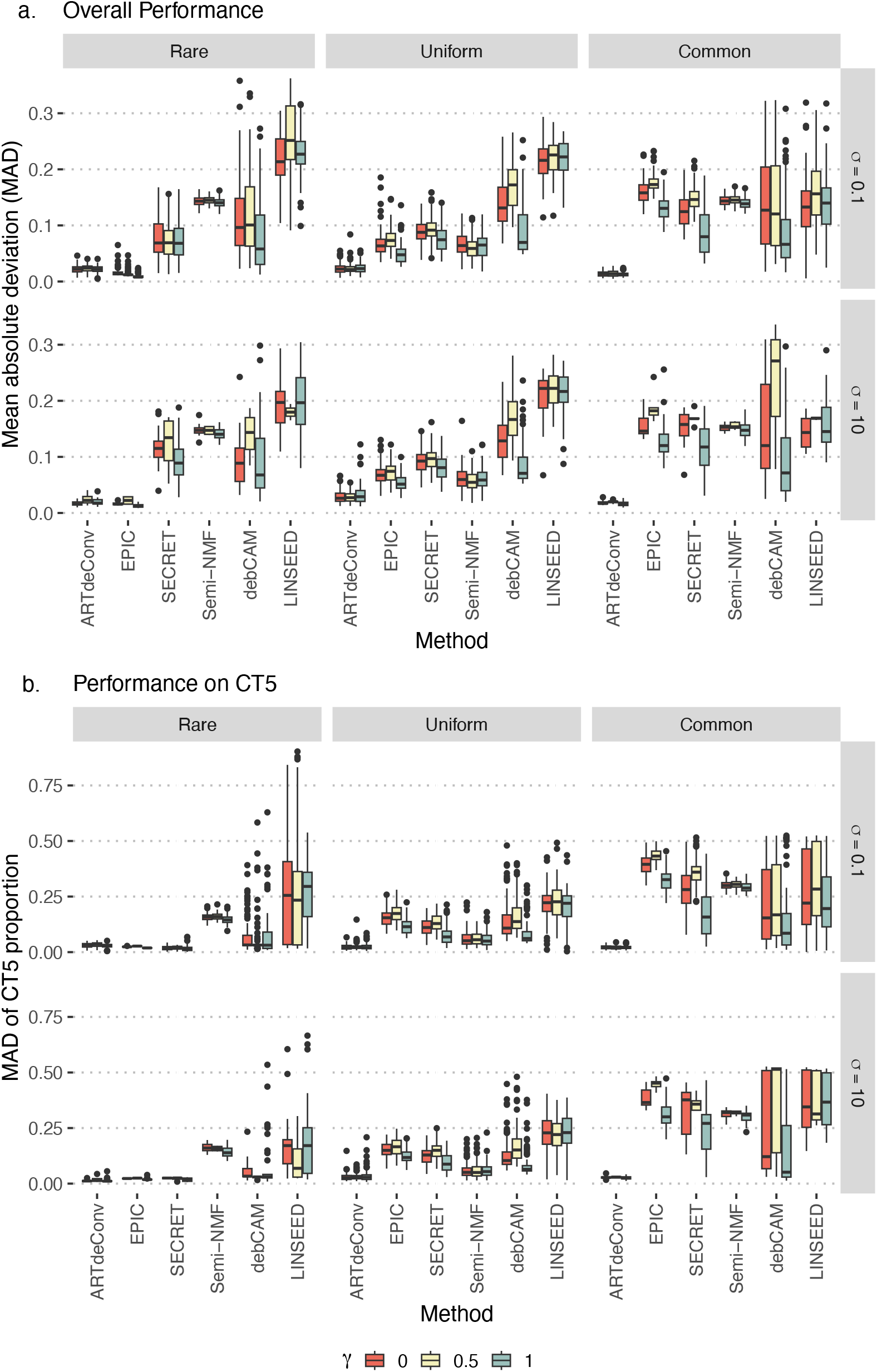
**a**. Mean absolute deviation (MAD) between true proportions and estimated cell-type proportions from simulated pseudo-bulks. **b**. MAD between estimated and true proportions of CT5 in the pseudo-bulks for the benchmark simulations. Each column represents a case of the relative abundance of CT5 against other cell types in the simulated pseudo-bulks. Each row represents the level of noises controlled by *σ*. Colors represent different levels of cell-type specificity of genes in **Θ**^*^ regulated by different *γ*.

Among the semi-reference-based methods, EPIC achieved high overall accuracy when CT5 was relatively rare in the pseudo-bulk samples. However, its performance in estimating the proportion of CT5 alone was surpassed by SECRET (Figure 2). While the overall performance of EPIC was comparable to that of SECRET when CT5 had similar abundance to other cell types, EPIC’s accuracy diminished when CT5 was relatively common among the pseudo-bulks. This is consistent with previous findings that EPIC tends to underestimate the proportions of cell types not characterized in the reference (Tai et al., 2021). Although SECRET showed higher accuracy than EPIC in scenarios where CT5 was common, its overall performance remained inferior to ARTdeConv and did not significantly enhance the accuracy of CT5 estimates over EPIC, except when CT5 was relatively rare, where it performed slightly better than ARTdeConv (Figures 2 and 3).

Among the reference-free methods, debCAM demonstrated accuracy that exceeded those of semi-reference-based methods when CT5 was relatively common (Figures 2 and 3). As expected, debCAM’s performance improved with highly cell-type-specific signature genes (*γ* = 1). Conversely, Semi-NMF performed well when CT5 had similar abundance to other cell types in the pseudo-bulk samples. LINSEED, known for its insensitivity to cell-type mRNA amounts, consistently showed the worst performance among the methods, except in a few specific cases.

In conclusion, ARTdeConv showed superior performance to its semi-reference-based peers and reference-free methods in our simulations. We further remark that, in the simulations described above, **Θ**^*^ included knowledge of marker genes for CT5. In practice, obtaining the marker genes of cell types without reference expression is, while not as straightforward, achievable through methods such as in Wang et al. (2016) and Qiu et al. (2021). Furthermore, even without well defined marker genes for those cell types, deconvolution by ARTdeConv is still completely feasible, for instance the analyses in Sections 3.2 and 3.3.

### 3.2. ARTdeConv Accurately Estimates Cell Type Proportions of PBMC from Bulk Gene Expression in a Human Influenza Vaccine Study

To benchmark the performance of ARTdeConv on real-world data and exemplify its application, we utilized a dataset sourced from Hoek et al. (2015). In this study, blood samples were obtained from healthy volunteer subjects who were administered a trivalent inactivated influenza vaccine (TIV). Both bulk PBMC samples and sorted PBMC cell lines from two enrolled subjects (“HD30” and “HD31”) vaccinated with a single dose of 2011–2012 seasonal TIV were collected at four different time points: before vaccination (Day 0), and on Day 1, 3, and 7 post-vaccination.

Deconvolution was performed on eight bulk PBMC samples whose gene expression were measured in transcript per million (TPM). Gene expression in TPM from the sorted PBMC cell lines of the two subjects on Day 0 were used to construct the partial signature matrix. The study characterized four distinct PBMC cell types: T cells, B cells, natural killer (NK) cells, and monocytes. Additionally, there were other unspecified cell types collectively termed into one super cell type as “others”, which could include, for instance, various types of dendritic cells and low density neutrophils (Kleiveland, 2015; Monaco et al., 2019). Detailed data processing steps could be referred to at Supplementary Material Section E.1.

The means and ranges of the cell-types in question were obtained through the reference values in Kleiveland (2015). For each cell type, the proportion means were calculated by taking the average of the upper and lower bounds of the reference values, while the ranges were calculated by taking the difference between the two bounds. For the “others” cell type, the mean was calculated by subtracting the means of all other cell types from one, and the range the difference between the upper and lower possible proportions of the other cell types. The exact values of the means and ranges are reported in Supplementary Material Table 1 (details of the calculation are given in Supplementary Material Section E.2). The vector of ranges was further normalized such that it has unit *L*_2_ norm before deconvolution. Additional details regarding the tuning grid and algorithm parameter setup of this analysis can be found at Supplementary Material Section E.3.

**Table 1:**
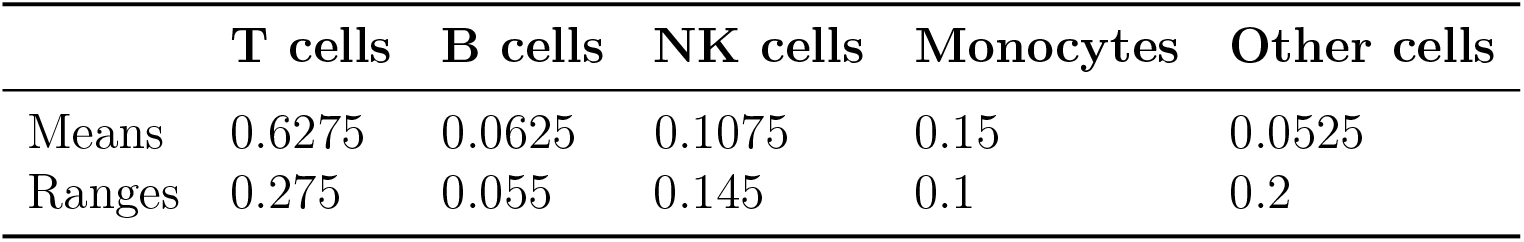
Means and ranges of PBMC cell type proportions calculated from the reference values.

The PBMC percentages on Day 0 were measured by flow cytometry (Hoek et al., 2015; Racle et al., 2017), which were used as the ground truths for evaluating the performance of ARTdeConv. Indeed, ARTdeConv achieved a notable degree of precision in estimating cell-type proportions on Day 0 as compared to the flow cytometry measurements (Figure 4a), achieving a canonical correlation coefficient (CCC) of 0.974 (Pearson’s correlation = 0.974, MAD = 0.036). This performance is on par with that of EPIC and surpassed those of CIBERSORT etc. as reported by Racle et al. (2017).

**Figure 4:**
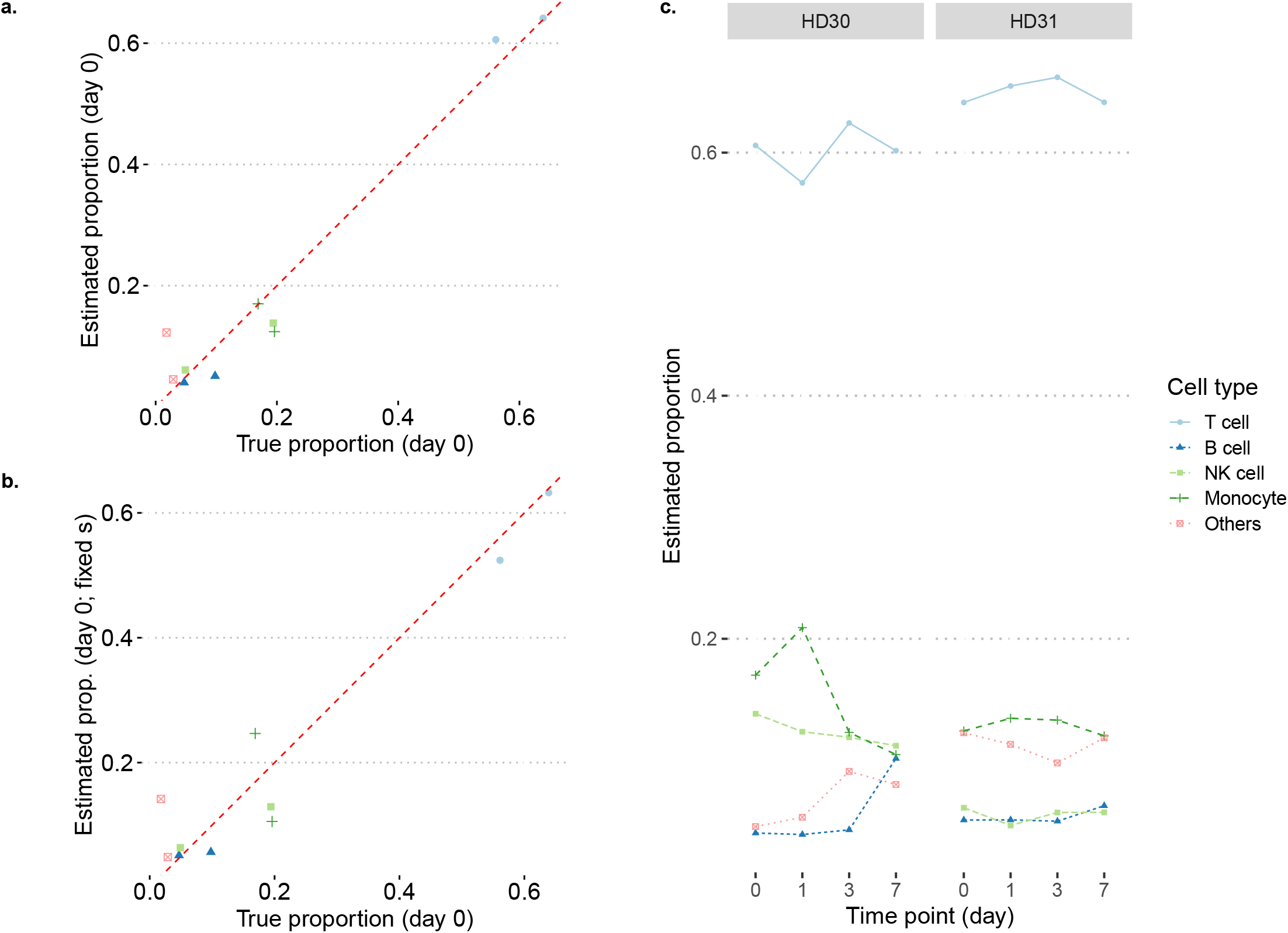
**a**. Estimated PBMC proportions by ARTdeConv versus true proportions measured by flow cytometry for two PBMC samples collected on Day 0 with flexible mRNA amount parameters. **b**. Estimated PBMC proportions by ARTdeConv versus true proportions measured by flow cytometry for two PBMC samples collected on Day 0 with mRNA amount parameters coerced to 1 for all cell types. **c**. Estimated PBMC proportions by ARTdeConv on all of the eight PBMC samples from two subjects across time points with flexible mRNA amount parameters.

We also estimated proportions of PBMC cell types across all time points utilizing the same partial signature matrix (Figure 4c). On HD30, a decline of T cell abundances before Day 1 and an increase between Days 1 and 3 were observed, followed by a slight decline until Day 7. An increase of monocyte abundances before and a decrease after Day 1 were also seen. Both trends are consistent with the profile of a virus shedder of the H1N1 virus in Rahil et al. (2020) with a small time shift. The time shift could be attributed to differences in viral strains and strengths between Rahil et al. (2020) and Hoek et al. (2015), as the former is a study on real H1N1 patients. The lack of notable changes in the cell-type proportions of HD31 resembles more to the profile of a non-virus shedder. More information on the viral progression of these two subjects is needed to confirm these observations.

We wish to remark on the utility of adjusting for cell-type mRNA amounts in this case. It is known that normalization by TPM loses such information, which was corrected by ARTdeConv via diag(***ŝ***). When re-running ARTdeConv on the same samples with diag(***s***) forced to be the identity matrix, we observed less accurate estimates of proportions (Figure 4b), corroborating the need for the adjsuetment in this scenario.

### 3.3. ARTdeConv Reveals Changes in Key PBMC Cell Type Proportion in COVID-19 Patients

We performed an extensive deconvolution analysis on PBMC bulk samples gathered from 17 healthy controls and 14 patients with COVID-19 diagnosis recruited for a systemic immunity assessment against COVID-19 infections in humans by Arunachalam et al. (2020). The study also classified the COVID-19 severity of infected patients. Patients in the study were designated with three levels of severity: moderate, severe, and Intensity Care Unit-hospitalized (ICU). Details regarding the patient recruitment and the classification of COVID-19 severity are in Supplementary Material Section F.1.

Bulk RNA-seq data downloaded using NCBI GEO accession number GSE152418 were utilized to perform deconvolution for estimating the proportions of the four major PBMC cell types: T cell, B cell, natural killer (NK) cell, and monocyte. The authors also included single cell RNA-seq data on separate independent blood samples from five healthy controls and seven COVID-19 patients. Of these, all healthy and six COVID-19 infected subjects had matching bulk and single cell samples that passed quality control. The single cell data were downloaded using GEO accession GSE155673 and were used to construct the partial signature matrix. Complete descriptions of the bulk and single cell data, their quality control and pre-processing, as well as the creation of gene signature matrices can be found in Supplementary Material Section F.1.

Deconvolution was performed separately on samples from healthy controls and COVID-19 patients using bulk and signature matrices that matched the disease status. For each set of samples, we assumed knowledge on the reference expression of the four major PBMC cell types: T cell, B cell, NK cell, and monocyte. All other cell types’ reference expression were assumed unkown and they were grouped into the super cell type “Others”. Since obtaining the reference means and ranges of the cell types separately for healthy and diseased populations proved infeasible, we opted to use the same set of numbers in Supplementary Table 1 for running ARTdeConv. Detailed descriptions of the tuning grid and other ARTdeConv parameters can be found in Supplementary Material Section F.2.

The complete results of deconvolved cell type proportions from ARTdeConv are shown in Figure 5a (boxplots) as well as Supplementary Figures 1 and 2 (bar charts). We observed that in COVID-19 samples, the proportions of T cells were notably lower than in the healthy control samples (Figure 5a). Among the diseased samples, T cell depletion were found in several severe and ICU samples (Figure 5b). This could be explained by T lymphopenia, commonly observed on blood samples of COVID-19 patients with severe symptoms but less frequently on those from patients with mild to moderate symptoms, as a result of the immunological responses of T cells to COVID-19 (Chen and John Wherry, 2020). It was also observed that Severe or ICU samples exhibited higher monocyte abundances in PBMC (Figure 5c). Similar trends have also been observed by Zhang et al. (2022) in blood samples of patients with severe COVID-19.

**Figure 5:**
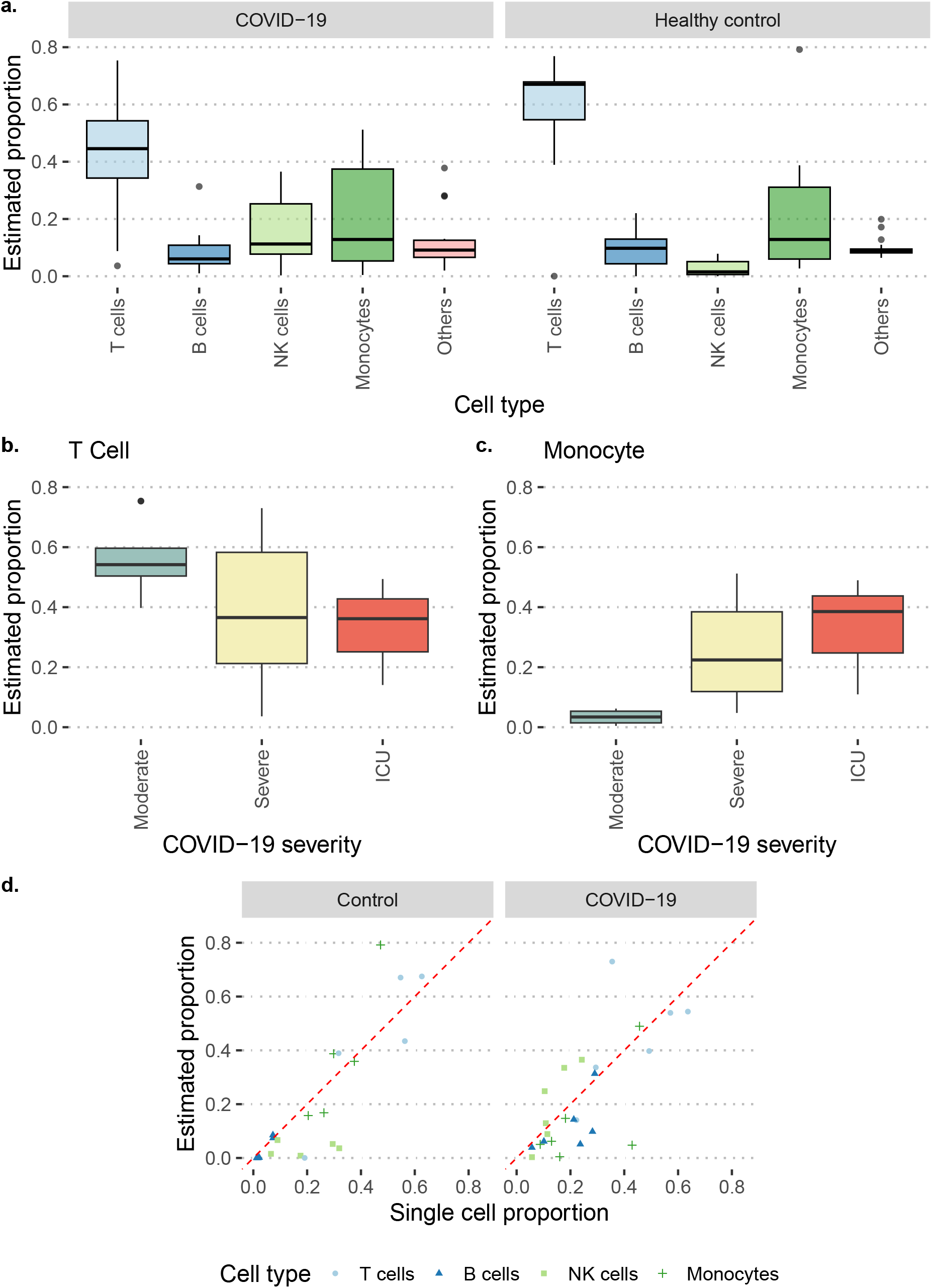
**a**. Box plots for estimated PBMC proportions by ARTdeConv in separate deconvolution analyses for healthy control and COVID-19-infected samples. **b**. and **c**. Box plots for estimated T cell and monocyte proportions on COVID-19-infected samples of different severity. **d**. Scatter plots for estimated PBMC proportions vs. matching tissue PBMC proportions from independent single-cell studies from five healthy controls and six COVID-19-diagnosed subjects.

Arunachalam et al. (2020) used the abundances of cells from the single-cell samples as a surrogate measurement for true abundances (except for dendritic cells, which were manually enriched in the single-cell samples). The relative deconvolved abundances of T cells, B cells, NK cells, and monocytes were then compared against the relative abundances of single cells. ARTdeConv demonstrated satisfactory accuracy in deconvolving cell-type abundances (Figure 5d), achieving a CCC of 0.815 among healthy control samples (Pearson’s correlation = 0.860, MAD = 0.098) and 0.694 among COVID-19 samples (Pearson’s correlation = 0.717, MAD = 0.103).

While we observed a slightly lower accuracy among COVID-19 patients compared to healthy controls, the COVID-19 patients spanned several disease severity classes and various duration of infection. These factors might increase the variation of gene expression, both on the cell type and the bulk levels, making deconvolution more challenging. The analysis also identified one outlier in healthy controls, S066 (Supplementary Figure 2). Given the typical high abundances of T cells in human PBMC samples, it was surprising that ARTdeConv did not detect any T cell in this sample, while attributed monocytes as the most abundant cell type. After further explorations in the expression of *CD3* and *CD14* genes, which are experimentally confirmed marker genes of T cells and classical monocytes respectively (Kleiveland, 2015), we discovered exceptionally low bulk *CD3* expression (Supplementary Figure 3) and high *CD14* expression (Supplementary Figure 4) in the sample from S066, which were consistent with this result.

## 4. Discussion

In this paper, we introduce ARTdeConv, an innovative deconvolution approach. An important feature of ARTdeConv is its adoption of a tri-factor model, which integrates the cell-type mRNA amounts during the deconvolution process. As a semi-reference-based method, ARTdeConv offers enhanced flexibility compared to reference-based methods, as it accommodates cell types whose reference gene expression is not known by grouping them into one super cell type, while presenting advantages over reference-free methods by incorporating the partial signature matrix. Moreover, the method makes effective use of reference information on proportion means and ranges derived from external studies.

Additionally, we derive the MU algorithm for ARTdeConv and present a theorem that establishes the convergence of this algorithm to stationary points. This proof is derived by casting ARTdeConv’s algorithm as a special case of the BSUM algorithm introduced by Razaviyayn et al. (2013).

On simulated pseudo-bulks, we demonstrated the advantages of ARTdeConv over other semi-reference-based methods. Notably, ARTdeConv performed better compared to EPIC when the cell types without reference expression became relatively abundant and had an overall edge against SECRET and other well-known reference-free methods. Moreover, both EPIC and SECRET require manual re-normalization of the estimated proportions by relatively precise cell-type mRNA amounts, specifications that are not required for ARTdeConv. In practice, such precise amounts can be difficult to obtain when the cell types with missing reference expression become numerous. Compared to quanTIseq and BayICE, ARTdeConv can flexibly utilize customized partial signature matrices, which are common in deconvolution applications. To the best of our knowledge, current semi-reference-based methods only take in one cell type or super cell type whose reference expression is unknown. A direction for future research in methodology is developing novel semi-reference-based methods that can distinguish multiple cell types with missing reference, i.e. truly allowing *K > K*_0_ + 1.

An argument can be made that obtaining precise reference information on proportions demands additional efforts in practice. However, the advantages of integrating this information are substantial. Without it, the task of associating estimated proportions with their respective cell types can prove challenging. First, the estimated proportions of the cell types without reference expression could be biased towards zero, as in the case of semi-reference-based methods like EPIC. Secondly, for all reference-free methods, we had to perform manual matching of results (as demonstrated in Section 3.1), which becomes unfeasible if multiple cell types lack reference expression.

On the algorithmic side, ARTdeConv provides a guarantee of convergence, closing a gap in the theoretical characterization of deconvolution methods in previous studies. Of note, ARTdeConv does not assure that the estimated proportions are globally optimal. This challenge is not particular to ARTdeConv and is shared by other deconvolution methods with iterative optimization procedures such as SECRET, NMF, and debCAM. To counter this, we recommend employing marker genes for the deconvolution process and considering multiple restarts of ARTdeConv. While methods like debCAM mitigate this issue by using a different mathematical framework, their assumptions are usually breached in practice, and their performance diminishes as a result.

We demonstrated the application of ARTdeConv to two different sets of real data. While the data from Hoek et al. (2015) had limited sample size, we validated ARTdeConv’s performance using the data from the more extensive study of Arunachalam et al. (2020). We remark that both sets of data offer an edge where both the bulk samples and the independent samples for making partial signature matrices were collected from independent samples concurrently, minimizing the influence of most technical artifacts. In situations where bulk and cell-type reference expression are derived from different studies, additional caution against those technical artifacts should be exercised during data pre-processing prior to deconvolution.

## Software Availability

ARTdeConv is available as an R package for download from the GitHub repository: https://github.com/gr8lawrence/ARTDeConv.

## Data Availability

Raw sequencing data in the fastq format for the analysis in Section 3.2 can be accessed through Sequence Read Archive (SRA) BioProject PRJNA271578. Measured proportions for PBMC cell types on Day 0 samples can be found in the supplementary materials of (Racle et al., 2017). Bulk and single cell data for the analysis in Section 3.3 could be accessed from NCBI GEO using the accession number GSE152418 and GSE155673 respectively.

## Acknowledgements

The authors gratefully acknowledge the support of National Institute of Allergy and Infectious Diseases at the National Institutes of Health through the grant U19AI144181 and that of National Institute on Aging through the grant R01AG073259.

## Supplementary Material

### A. Technical Assumptions

Before presenting the main results, we state the technical assumptions.

#### Assumption 1

(Initial conditions). *(i)* ∀*k*, ∃*j*_*k*_ ∈ {1, …, *m*} *such that* 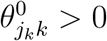*; (ii)* ∀*k*, ∃*i*_*k*_ ∈ {1, …, *n*} *such that* 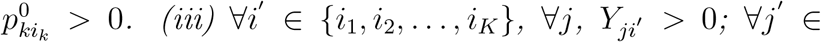 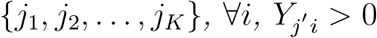.

#### Assumption 2

(Boundedness). *(i) For any* 1 ≤ *j* ≤ *m and* 1 ≤ *i* ≤ *n, Y*_*ji*_ ≤ *M*_*Y*_ *<* ∞. *(ii) For any* 1 ≤ *j* ≤ *m and* 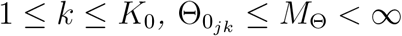.

These assumptions are reasonable in practice. For every cell type with established reference expression, it is safe to assume that there exist some genes that display non-zero cell-type-specific expression, thereby fulfilling Assumption 1(*i*). For cell types lacking reference expression, an initial value in **Θ**^0^ can be straightforwardly set to a positive value, thereby also satisfying the same assumption. Assumption 1(*ii*) is automatically satisfied due to how ***P*** ^0^ is initialized in Algorithm 1. Assumption 1(*iii*) plays a pivotal role in ensuring that positive initialized values remain positive throughout the iterative process (see Supplementary Material Section B for details). This assumption is easy to verify and holds true unless the bulk expression matrix is excessively sparse, which is uncommon in practice. Assumption 2 is a mild assumption on upper bounds of gene expression.

### B. The MU Steps

#### B.1. A Preliminary Lemma

Similar to Lee and Seung (2000), *f* (**Θ, *s, P***) is minimized using block-wise auxiliary functions that are quadratic over-estimators, which then leads to the MU steps in (4), (5), and (6). The claims that these updates will keep the positive initialized values positive will be investigated at the end of this section. An auxiliary function is defined next.

##### Definition 1

(Auxiliary function). *Given a vector space V and* ***v, v***^*′*^ ∈ *V, a function g*(***v***|***v***^*′*^) : *V* × *V* ↦ ℝ *is an auxiliary function for the function f* (***v***) : *V* ↦ ℝ *at* ***v***^*′*^ *if the following conditions are satisfied:*

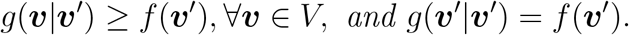

As we shall see, the vector blocks in the auxiliary functions would be each row of **Θ**, each column of ***P***, and ***s***. To find the specific auxiliary functions for *f* (**Θ, *s, P***), a slightly modified form of a technical lemma given by Lin (2007) is needed and stated next. Its proof is also given in the same paper and thus omitted.

##### Lemma 1.

*Given a positive semi-definite matrix* 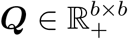 *and* 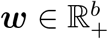, *let I* ⊆ {1, …, *b*} *be a set of indices such that w*_*k*_ *>* 0 *for any k* ∈ *I*, 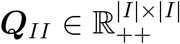, *and* 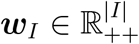 *be the sub-matrix of* ***Q*** *and the sub-vector of* ***w*** *with indices in I, where* |*I*| *denotes the cardinality of I. Then*,

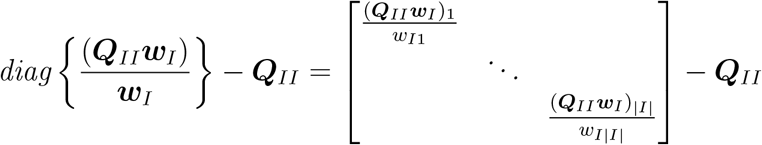

*is always positive semi-definite*.

With Lemma 1, the auxiliary functions and multiplicative update (MU) steps are ready to be derived.

#### B.2. Derivations of The MU Steps

In addition to those in Section 2.1, we introduce two additional notation: for two square matrices of the same dimension, we write ***A*** ≻***B*** and ***A*** ⪰ ***B*** if ***A*** − ***B*** is positive-definite and positive-semidefinite, respectively.

The auxiliary function and update step for each row 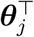 is derived and then combined for *j* = 1, …, *m* into the form of (4). For simplicity in notations, the index *j* is omitted in subscripts. The iteration number *t* is also omitted for ***s*** and ***P*** since both are treated as fixed. That is, we denote 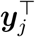 as ***y***^⊤^, 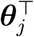 as ***θ***^⊤^, and 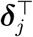 as ***δ***. We also define 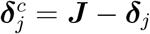 for all *j*. Thus, for the *j*-th row of **Θ**, we can define the sub-problem of *f* (**Θ, *s, P***) as

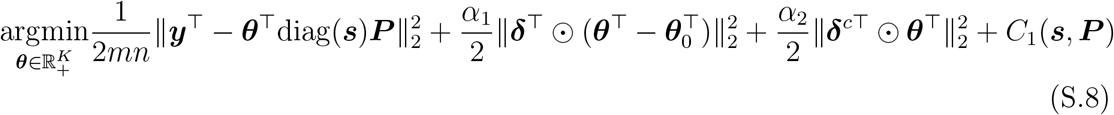

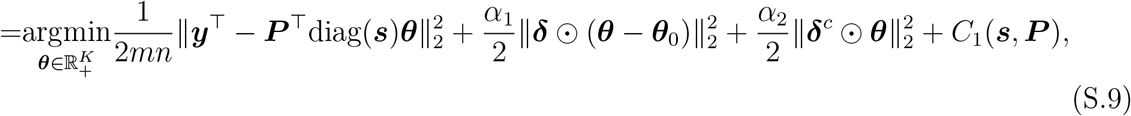

where diag(***s***) and ***P*** are treated as fixed and *C*_1_(***s, P***) is a constant with respect to ***θ***. We call the function in (S.9) *f*_*θ*_(***θ***). (S.9) is derived from (S.8) by transposing. Now, to find the auxiliary function *h*_*θ*_(***θ***|***θ***^*t*^), by performing a second-order Taylor series expansion on *f*_*θ*_(***θ***) at ***θ*** = ***θ***^*t*^, we have

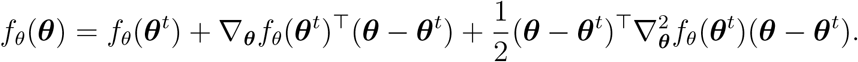

Next, letting ***V*** (***δ***) = diag(***δ***) and ***V*** ^*c*^(***δ***^*c*^) = diag(***δ***^*c*^), it can be observed that ***δ*** ⊙ ***θ*** = ***V θ*** and ***δ***^*c*^ ⊙ ***θ*** = ***V*** ^*c*^***θ***. Also, ***V*** + ***V*** ^*c*^ = ***I***. The first and second derivatives of *f*_*θ*_(***θ***) are thus:

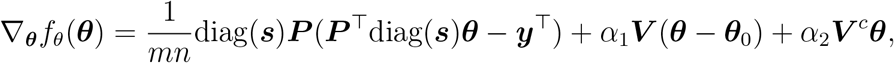

and

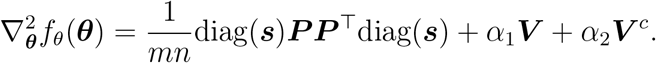

It can be guaranteed 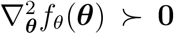 given *α*_1_, *α*_2_ *>* 0, as 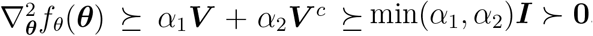. Then, define 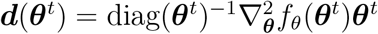, and ***D***(***θ***^*t*^) = diag(***d***(***θ***^*t*^)). Lemma 1 implies that, on coordinates that correspond to the positive elements of ***θ***, 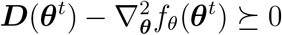. Define 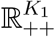, where *K*_1_ ≤ *K*, as the open half space that contains this positive subset of ***θ***. For simplification, we use the same notations for the full vectors and matrices on those positive coordinates. As mentioned in Section 2.2, we shall see that once a coordinate of ***θ*** reaches zero, it stays zero thereafter, which justifies the focus on these positive coordinates. Next, define

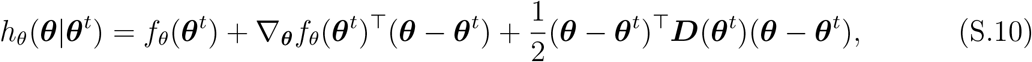

which is an auxiliary function of (S.9) (thus of *f* (**Θ, *s, P***) when we combine all sub-problems together for *j* = 1, … *m*). To demonstrate that it satisfies Definition 1, simply see that

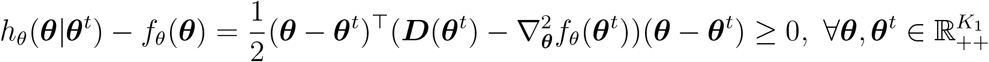

and due to Lemma 1 and that *h*_*θ*_(***θ***^*t*^|***θ***^*t*^) − *f*_*θ*_(***θ***^*t*^) = 0,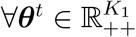. To derive the MU step from this auxiliary function, we find ***θ***^*t*+1^ such that

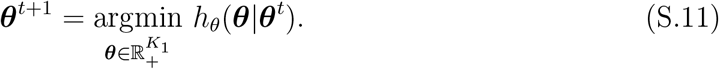

To guarantee the existence of a solution for (S.11), the feasible set of ***θ*** is a closed space that includes 0. Since 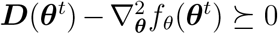 and 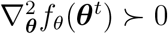 by the assumptions, ***D***(***θ***^*t*^) ≻ 0, which implies that (S.10) is strictly convex. Thus, (S.10) has a global minimizer in the closed half space 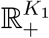. By the optimality condition, if a feasible 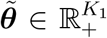 satisfies 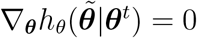, then 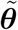 minimizes (S.10) globally and 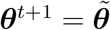 becasue of (S.11). Thus, we solve

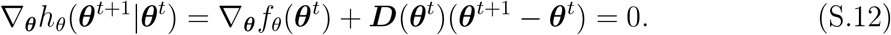

Because ***D***(***θ***^*t*^) is diagonal, for each *k* ∈ 1, …, *K*_1_, (S.12) implies

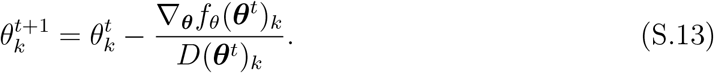

We further simplify the notations by letting

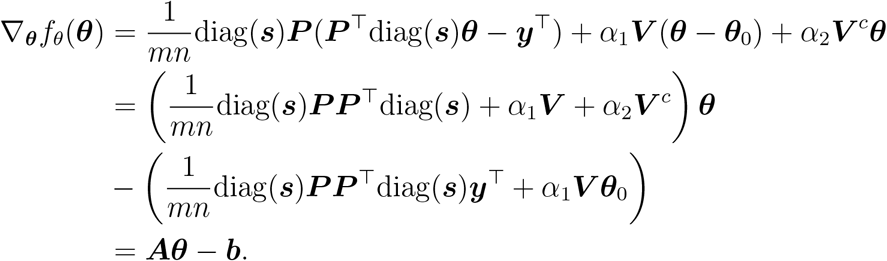

It follows immediately that 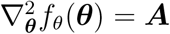. Plugging in the values of ∇_***θ***_*f*_*θ*_(***θ***^*t*^)_*k*_ and *D*(***θ***^*t*^)_*k*_ to (S.13), we have

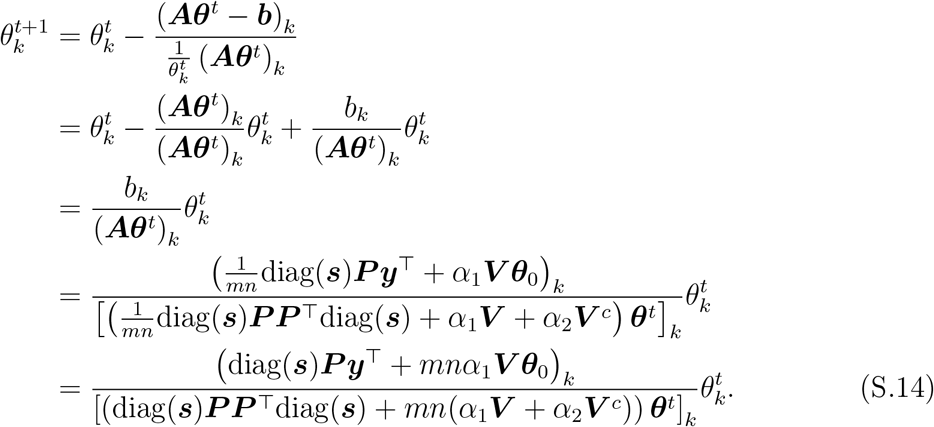

The multiplicative form of (S.14) guarantees that ***θ***^*t*+1^ is feasible. Therefore, ***θ***^*t*+1^ satisfies (S.11).

It is straightforward to check that (S.14) also applies to zero coordinates of 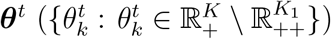, as zero coordinates remain zero after any multiplication. Therefore, it is the unified update step for the entire 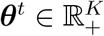. Since it corresponds to the update for the (*j, k*)-th element in **Θ**, combining all elements gives (4).

We remark that (S.13) reflects the block coordinate descent nature of the multiplicative update in **Θ**, as observed by Kim et al. (2014). In fact, the update steps for ***P*** and diag(***s***) can be written in similar fashions.

The update step for ***P*** in (5) can be similarly derived as that for **Θ**. We begin with specifying the sub-problem for ***p***_*i*_, *i* = 1, …, *n*, treating **Θ** and diag(***s***) as fixed and omitting the subscripts and the iteration number (*t* for ***s*** and *t* + 1 for **Θ**) for convenience in notations:

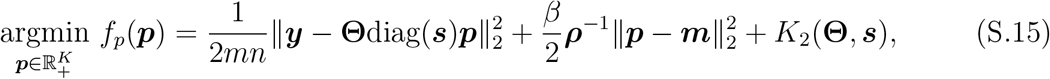

where *K*_2_(**Θ, *s***) only depends on **Θ** and ***s*** and is a constant with respect to ***P***. Then, its first and second derivatives with respect to ***p*** are

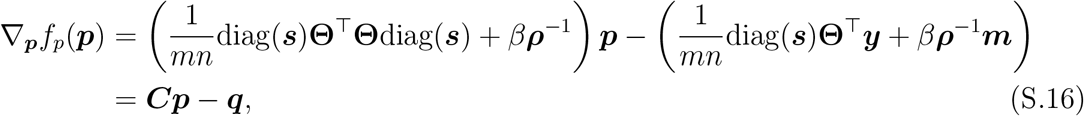

and

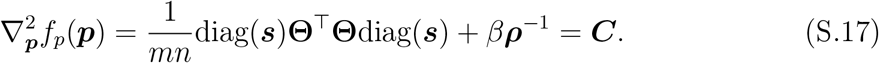

To find the auxiliary function of (S.15), define 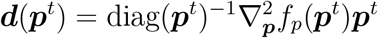, and ***D***(***p***^*t*^) = diag(***d***(***p***^*t*^)), and

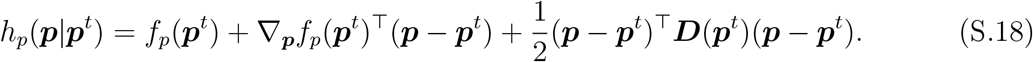

On the *K*_2_ ≤ *K* positive coordinates in ***p***^*t*^, (S.18) is the auxiliary function of (S.15) due to Lemma 1. The verification is the same to that for *h*_*θ*_(***θ θ***^*t*^) and omitted here. Next, we find ***p***^*t*+1^ such that

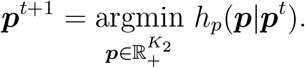

This in turn corresponds to finding ***p***^*t*+1^ such that

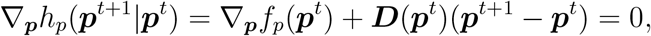

given that such a ***p***^*t*+1^ is feasible. After some algebra, for any *k* = 1, …, *K*_2_,

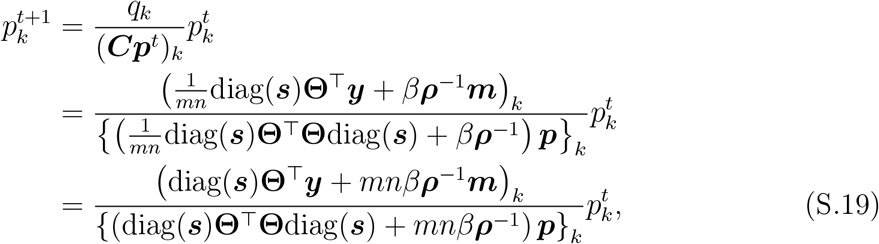

which produces a feasible ***p***^*t*+1^. Moreover, (S.19) also applies to the zero coordinates in ***p***^*t*^. Since (S.19) corresponds to the update step for the (*k, i*)-th element for the ***P*** matrix, combining all of these elements yields (5).

Deriving the updates step for diag(***s***) in (6) largely follows the same footsteps as for **Θ** and ***P***. To find a suitable sub-problem, we start from the following identity

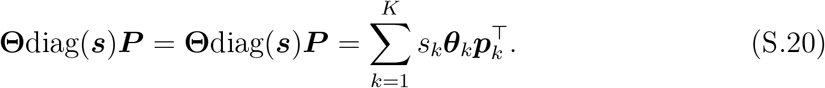

Denoting 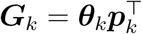 and only assuming ***s*** a variable, we transform the objective function in *f* (**Θ, *s, P***) with respect to ***s*** as follows

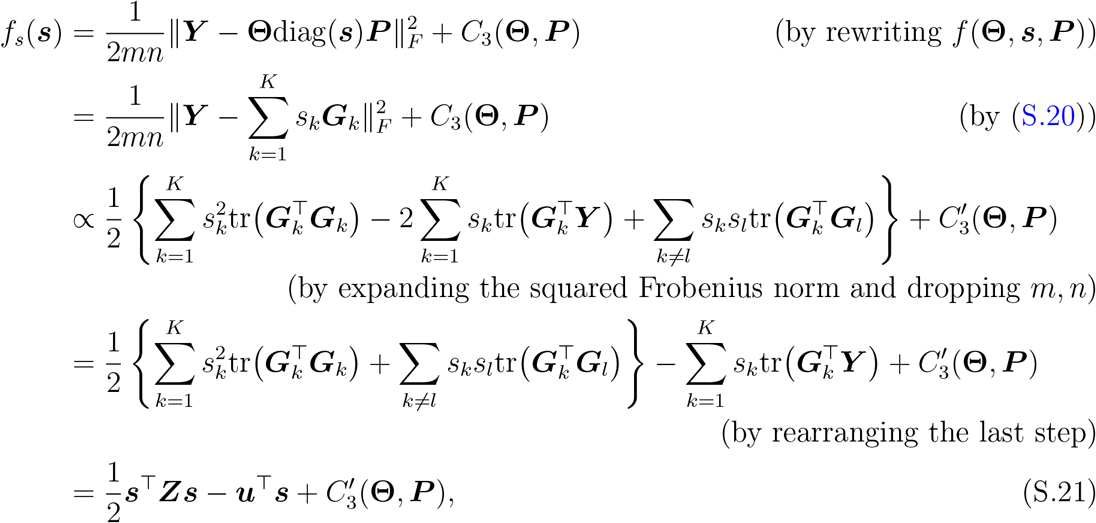

where 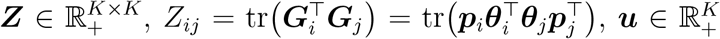, and 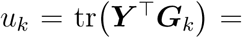 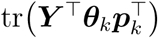. Both *C*_3_ and 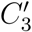 only depend on **Θ** and ***P***, hence are constant with respect to ***s***. (S.21) is then our sub-problem for ***s***. From (S.21), the first and second derivatives for ***s*** are ∇_***s***_*f*_*s*_(***s***) = ***Zs*** − ***u***, and 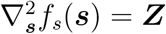. Again, define 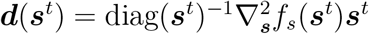, ***D***(***s***^*t*^) = diag(***d***(***s***^*t*^)), and

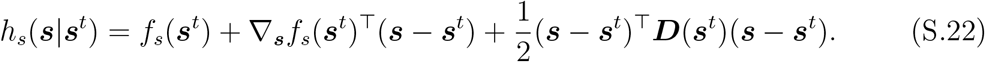

(S.22) is an auxiliary function of *f* (**Θ, *s, P***) for ***s*** according to Lemma 1. Since 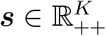, there is no need for sub-setting to positive coordinates. Thus, we obtain ***s***^*t*+1^ by finding a feasible

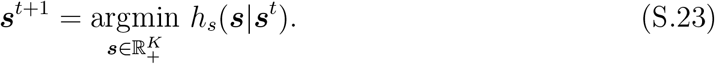

We remark that this is a slight relaxation of the problem to update ***s***, since it is originally assumed that 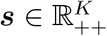. However, this allows us to obtain a guaranteed closed form global minimizer for (S.23) in the closed half space 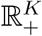, and the resulting update satisfies is feasible in 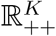 as long as not all values of ***θ***_*k*_ or 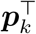 are 0 for any *k* = 1, …, *K*, which is stated as Assumption 1(*i*) and 1(*ii*). (S.23) leads to solving

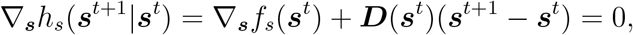

from which we have

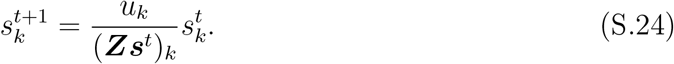

(S.24) corresponds to the update for the *k*-th coordinate in (6), which yields feasible 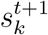 under the assumptions, since 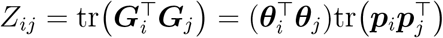 and 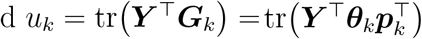. Moreover,

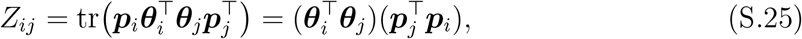

and that

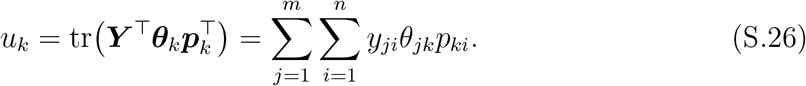

For the denominator in (S.24), we see that

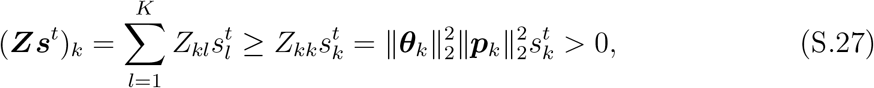

where the last step is a result of (S.25) and the stability of updates based Assumption 1(*i*) and 1(*ii*), which will be discussed in details below. As for the numerator of (S.24), taking from the stability of updates due to Assumption 1(*i*) and 1(*ii*) and assuming 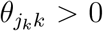 and 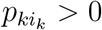, by (S.26), we have

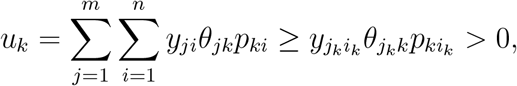

as long as 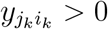. Given Assumption 1(*iii*), we conclude that 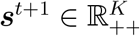.

#### B.3. Stability of The Update Steps

To justify the claim that for any *j* and *k*, 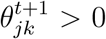 whenever 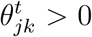 for *t* = 0, 1, 2, …, notice that the denominator of (S.14) is automatically positive whenever *α*_1_, *α*_2_ *>* 0. For guaranteeing that its numerator is also positive, we want to show

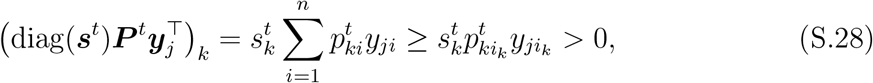

where the index *i*_*k*_ is defined as in Assumption 1(*ii*). By Assumption 1(*iii*), 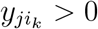, and 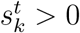 as discussed above. Thus, (S.28) holds whenever 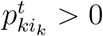. Similarly, for proving the claim that for any *k* and *i*, 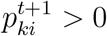 whenever 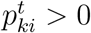, notice that the denominator for (S.19) is also positive whenever *β >* 0. To prove that its numerator is positive, we need to show

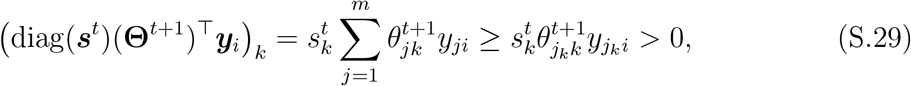

where *j*_*k*_ is defined as in Assumption 1(*i*). (S.29) holds whenever 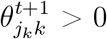.

Since by Assumption 1(*ii*) and Assumption 1(*i*), 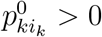 and 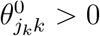. Therefore, we can show that 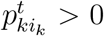 and 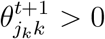 for any *t* = 0, 1, 2, … by induction through (S.28) and (S.29). Thus, the claims on the stability of updates hold.

### C. Proof of Theorem 1

To prove Theorem 1, we leverage on the convergence properties of the block successive upper-bound minimization (BSUM) algorithms proved by Razaviyayn et al. (2013). Namely, we can view Algorithm 1 as a BSUM algorithm by counting a total of *m* + *n* + 1 separate blocks which the algorithm minimizes successively: *m* for updating **Θ**^*t*^, *n* for ***P***^*t*^, and one for ***s***^*t*^. To utilize such properties, we need to prove the following proposition, which states the conditions on the update steps’ auxiliary functions for Algorithm 1 to formally qualify as BSUM, which is translated from Assumption 2 of Razaviyayn et al. (2013):

#### Proposition 1

(Algorithm 1 is BSUM). 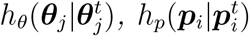, *and h*_*s*_(***s***|***s***^*t*^) *satisfy the following conditions for any j* = 1, 2, …, *m and i* = 1, 2, …, *n: (1)* 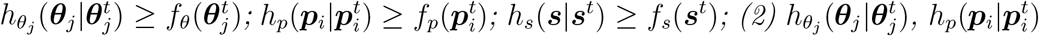, *and h*_*s*_(***s***|***s***^*t*^) *are continuous on their domains; (3) Any directional derivative of* 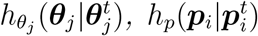, *and h*_*s*_(***s***|***s***^*t*^) *within their domains are equal to the direactional derivative of* 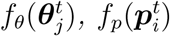, *and f*_*s*_(***s***^*t*^) *respectively*.

*Proof*. The proof is quite straightforward. (1) and (2) are direct from the construction of the surrogate functions and Lemma 1, which also manifests the majorization-minimization (MM) property (Lange, 2016) of each block’s update. (3) is seen by differentiating (S.10), (S.18), and (S.22), which leads to 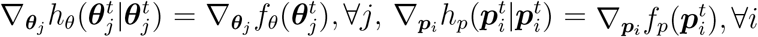, and ∇_***s***_*h*_*s*_(***s***^*t*^|***s***^*t*^) = ∇_***s***_*f*_*s*_(***s***^*t*^). Equal gradients automatically guarantee equal directional derivatives, proving the proposition.

Now that Algorithm 1 is found to be BSUM, Theorem 1 can be proved directly by the results of Theorem 2(b) in (Razaviyayn et al., 2013). But a few additional conditions presented in the following propositions need to be verified, both of which are proved thereafter.

#### Proposition 2

(Compact Sub-level Sets). *The sub-level set χ* ^0^ : {(**Θ, *s, P***) : *f* (**Θ, *s, P***) ≤ *f* (**Θ**^0^, ***s***^0^, ***P*** ^0^)} *of f* (**Θ, *s, P***) *given the update steps in Algorithm 1 is compact*.

#### Proposition 3

(Unique Global Minimizer of Auxiliary Functions). *Given Assumption 1*, ∀*t* ∈ ℝ _+_ *and for all i and j, the auxiliary functions* 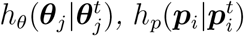, *and h*_*s*_(***s***|***s***^*t*^) *each have a unique global minimizer in* 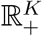.

#### C.1. Proof of Proposition 2

We first introduce the definition of coercive functions through:

##### Definition 2

(Coercivity). *A function f* (***X***_1_, …, ***X***_*N*_) : *V*_1_× … ×*V*_*N*_ → ℝ, *where V*_*i*_ *is a Euclidean vector space endorsed with the L*_2_ *norm if its elements are vectors or with the Frobenius norm if the elements are matrices, is called coercive if* ∀*i*, ∥***X***_*i*_∥ → ∞ *implies f* (***X***_1_, …, ***X***_*N*_) → ∞.

In fact, continuous coercive functions always have compact sub-level sets, as shown by the following lemma:

##### Lemma 2.

*If a continuous function f* (***X***_1_, …, ***X***_*N*_) : *V*_1_ × … × *V*_*N*_ → ℝ *is jointly coercive, then its sublevel set* 𝒳 ^*r*^ := {(***X***_1_, …, ***X***_*N*_) : *f* (***X***_1_, …, ***X***_*N*_) ≤ *r*} *is compact for any r* ∈ ℝ.

*Proof*. Because *f* is continuous and the space *f* (***X***_1_, …, ***X***_*n*_) ≤ *r* is closed, 𝒳 ^*r*^ is closed. We prove the boundedness of *χ*^*r*^ by contrapositive. We let 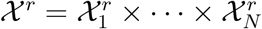, where 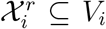. If *χ*^*r*^ is unbounded, then one of 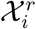 is unbounded. If 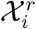 is unbounded, we can find a sequence of 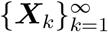 such that ∥***X***_*k*_∥ → ∞ as *k* → ∞, which implies *f* is not jointly coercive. Therefore, whenever *f* is jointly coercive, 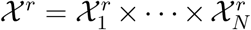 is bounded. Since Euclidean spaces admit the Heine-Borel property, *χ*^*r*^ is compact. □

Coming back to the proof of Proposition 2, it is easy to check that *f* (**Θ, *s, P***) is continuously differentiable. To prove that it is also coercive, we need to show that *f* (**Θ, *s, P***) tends to ∞ whenever ∥**Θ**∥_*F*_, ∥***s***∥_2_, or ∥***P***∥_*F*_ does so.

Beginning with the case for **Θ**, we have, by *f* (**Θ, *s, P***),

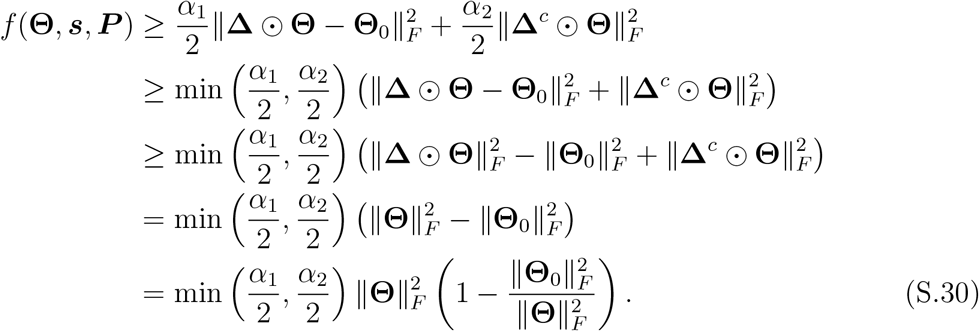

Since 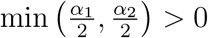 and 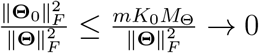 as ∥**Θ**∥_*F*_ → ∞ by Assumption 2, (S.30) then guarantees that *f* (**Θ, *s, P***) → ∞ when ∥**Θ**∥_*F*_ → ∞.

In the case for ***P***, whenever ∥***P*** ∥_*F*_ → ∞, there is a *k* such that 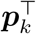 satisfies 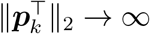. Also, since 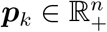, there exists a column indexed *i*_*k*_ as defined in Assumption 1(*ii*) such that 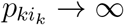. Then,

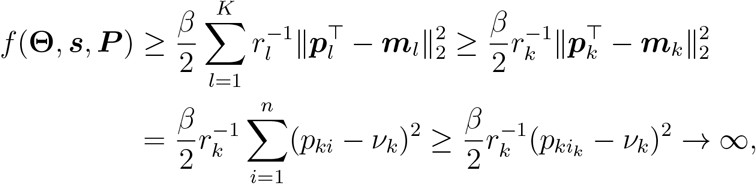

as *ν*_*k*_ is assumed fixed and *β >* 0.

Lastly for ***s***, we can similarly argue that, because 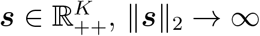 implies there is a *k* such that *s*_*k*_ → ∞. From the steps leading to (S.21), we have

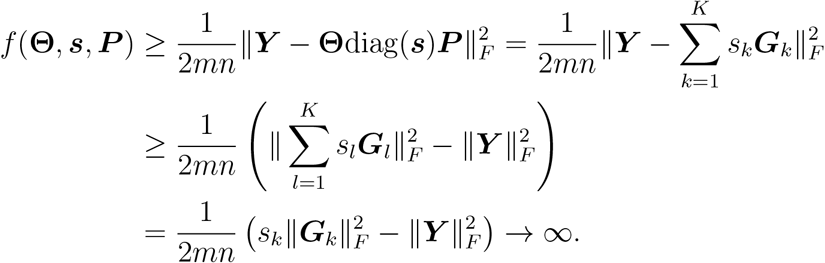

The last line is because ∥***G***_*k*_∥ *>* 0 due to Assumption 1(*i*) and Assumption 1(*ii*) and that ∥***Y***∥_*F*_ ≤ *mnM*_*Y*_ *<* ∞ by Assumption 2. Combining the cases for **Θ, *P***, and ***s*** completes the proof that *f* (**Θ, *s, P***) is coercive.

Since *f* (**Θ, *s, P***) is continuous and coercive, the set *χ*^0^ : {(**Θ, *s, P***) : *f* (**Θ, *s, P***) ≤ *f* (**Θ**^0^, ***s***^0^, ***P*** ^0^)}is compact according to Lemma 2. We have concluded the proof of Proposition 2.

#### C.2. Proof of Proposition 3

Since 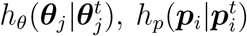, and *h*_*s*_(***s***|***s***^*t*^) are quadratic functions, proving Proposition 3 is equivalent to showing their Hessians are positive definite. Also, we only focus on the positive coordinates of 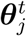 and 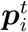 since other coordinates will remain zero through updates, and the proposition is vacuously true. By (S.10), (S.18), and (S.22), these Hessians are 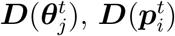, and ***D***(***s***^*t*^) respectively. Since they are all diagonal matrices, we want to show that their diagonal values are all positive.

For 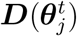, we have 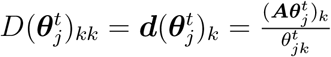. The denominator of 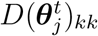 is positive. Dropping the index *j*, its numerator is

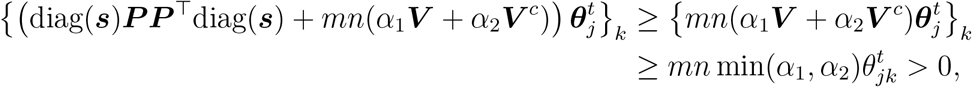

since ***V*** + ***V*** ^*c*^ = ***I***. Hence, 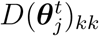 is positive.

For 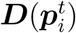, from (S.16) and (S.17), we also have 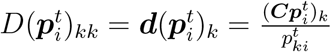. Likewise, the denominator is positive, and the numerator after dropping the index *i* is

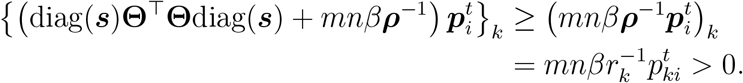

Lastly, *D*(***s***^*t*^)_*kk*_ is already shown to be positive from the conclusion of (S.27). This completes the proof of Proposition 3.

To finally prove Theorem 1, we combine the results of Propositions 1-3, and apply these conclusions to Theorem 2(b) in Razaviyayn et al. (2013). This way, we have shown that Algorithm 1 converges to a set of stationary points of *f* (**Θ, *s, P***).

### D. Supplementary Information for The Simulated Pseudo-Bulk Data

#### D.1. Data Generation

For *k* = 1, 2, …, 5, 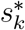 was generated independently from a *χ*^2^ distribution with 5 degrees of freedom. For the rare class, 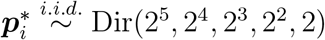, where Dir represents the Dirichlet distribution. For the uniform class, we first drew ***α***_*i*_ from a Dirichlet distribution with 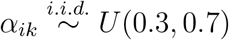, *k* = 1, …, 5, and then generated 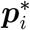 from Dir(***α***_*i*_). For the extra class, 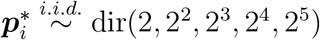, in reverse of the rare class. To generate values in **Θ**^*^, for each gene *j* = 1, 2, …, *M*, a 5 × 1 vector of expression was generated by

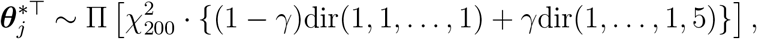

where Π(·) is the sampling without replacement operator. Π(·) allowed all of CT1-CT5 have a number of gene signatures.

The random errors ***ϵ*** were generated based on the principle of mean-variance dependency in gene expression data: the higher the mean of a gene’s expression, the higher their variation (Anders and Huber, 2010). For each gene *j*, its error-free mean of bulk expression 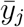 was calculated. Then its associated error for each sample *i* was generated by 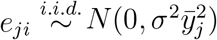, *i* = 1, 2, …, *n*,.

#### D.2. Method For Selecting Highly Cell-Type-Specific Genes

First, for each gene, the cell type for which it had the highest expression was picked as the target cell type. This gene then became a candidate for a marker of this target cell type. Second, for each candidate marker, the ratio of expression in its target cell type to those in the cell type with the second highest expression was calculated. Third, for each cell type, its candidate marker genes were ranked by the ratio calculated in the second step from highest to lowest. Finally, the top 100 candidate marker genes for each cell type were selected.

#### D.3. Tuning Grid And Algorithm Parameters

Cross-validation (CV) was performed in each simulation to find the best values for *α*_1_ and *β* according to Section 2.4. A tuning grid **𝔄**_1_ = **𝔅** = (10^−2^, 10^−1^, …, 10^3^, 10^4^) was set up with the tolerance parameter chosen as ***δ*** = 10^−5^.

#### D.4. Matching The Estimated Proportions to Cell Types For Reference-Free Methods

We utilized 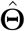 and the partial signature matrix 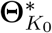. Starting from the first column of 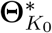, 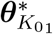, we found the column of 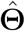, say column *k*, that had the highest Pearson’s correlation with 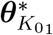. Then, the *k*-th row of 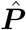 was determined to be estimated proportions for CT1.We did this for the rest of the columns of 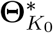 and those of 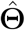, until there was one last unmatched column in 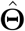. That unmatched column, and its corresponding row in 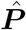, were then matched to CT5.

### E. Supplementary Method And Results For The Deconvolution of Bulk PBMC Samples from The Human Influenza Vaccine Study

#### E.1. Data Processing

FASTQ files containing 50bp (base pairs) pair-end raw nucleotide sequence reads of transcripts from Illumina Hiseq 2000 sequencers were downloaded from the Sequence Read Archive (SRA) BioProject PRJNA271578. The raw reads were pre-processed and filtered where reads with length < 50bp, with > 30% bases with quality scores < 30, having an average quality score < 25 in any 10bp interval, or corresponding to special adapter sequences were removed using the software `fastp` (, Chen et al., 2018). Then, reads passing the filters were mapped to reference human genome hg38 GRCh38 Release 43 (downloaded from the GENCODE website) and quantified using `salmon` (, Patro et al., 2017). The output files were then processed using the R package `tximport` (, Soneson et al., 2015): only protein-coding genes with an official gene symbol were selected and their transcripts per million (TPM) values were gathered into a gene-by-sample matrix of gene expression. Additionally, genes with a total TPM of less than 10 across the four cell types were excluded from consideration.

Distinct marker genes for the four cell types were individually selected for each subject using sorted bulk profiles of individual cell types. The selection process identified the top 10 genes exhibiting the highest cell-type specificity for each cell type from each subject according the ratio of expression method outlined in Section 3.1. The union of the two sets of marker genes, totalling 73 genes, were used in the deconvolution.

#### E.2. Calculating Reference PBMC Proportion Medians and Ranges from External Data

To calculate the reference PBMC cell-type proportion medians and ranges in Table 1, we adopted the data from Kleiveland (2015), a highly cited textbook on PBMC. The median and range for the cell types with reference expression were calculated through the descriptions in the textbook using a simple method. For example, the book states that lymphocytes account for 70% - 90% of all PBMCs, and among the lymphocytes 70% - 85% are T cells. Thus, the range of T cell percentage among PBMCs is 49% - 76.5%, or 0.49 - 0.765 in proportion. The median is thus (0.49 + 0.765)*/*2 = 0.6275 and the range is 0.765 − 0.49 = 0.275, corresponding to the reference values for T cell in Table 1. The reference values for B cell, NK cell, and monocyte were similarly calculated. For the “other cells”, their median proportion is 1 − (0.6275 + 0.0625 + 0.1075 + 0.15) = 0.0525. As for the ranges, the upper limit of the proportion for “other cells” occurs under the lower limit of lymphocytes and monocyte, which according to Kleiveland (2015) is 1 − (0.7 + 0.1) = 0.2 (70% lymphocyte, which includes T cell, B cell, and NK cell combined, and 10% monocyte), while the lower limit is 0 (when there are 90% lymphoyte and 20% monocyte, an impossible scenario). This leads to a range of 0.2.

We notice a range of 0.2 could be a little wide for “other cells”. As it turned out, these coarse parameters for medians and ranges sufficed for ARTdeConv to work properly. Thus, it is not always necessary to obtain super accurate reference parameters as long as they are in the right orders of magnitude.

#### E.3. Tuning Grid And Algorithm Parameters

A random seed of 1000 was set in the R programming environment. The tuning grid for *α*_1_ was set as the interval from 2^−5^ to 2^0^ with a step size of 0.2 in the power, and that for *β* was set as the interval from 2^0^ to 2^5^ also with a step size of 0.2 in the power. *α*_2_ was fixed at 10^−12^. The difference in magnitudes between *α*_1_ and *β* in the tuning grids ensured proper regularization, for there was a scale difference between gene expression and the proportions. A 4-fold cross-validation was invoked to choose the optimal tuning parameters. The eventual selected tuning parameter values were within the boundaries of the grids. The tolerance parameter was set as *δ* = 10^−4^.

#### E.4. Fixing mRNA Amounts as One For All Cell Types using ARTdeConv Results in Biased Deconvolution Results

In Section 3.2, it was posited that bulk and gene signature matrices whose expression values were measured in TPM would lose the information on cell-type mRNA amounts, necessitating the inclusion of ***s*** described in the underlying model (1). To verify this claim, we re-ran the data analysis on the same data in TPM as in Section 3.2 using the same empirical estimates ***M*** and ***ρ***, the same tolerance parameter, and the same tuning grid for the hyperparaters, and the same random seed, but coerced diag(***s***) to be the identity matrix (i.e. coerced the cell-type mRNA amounts to 1) throughout updates in Algorithm 1. We observed that the estimated proportions for PBMC cell types for both samples on Day 0 deviates more from those measured by flow cytometry when compared to when diag(***s***) is not coerced, as shown in Figure 4b.

### F. Supplementary Method And Results For The Deconvolution of Bulk PBMC Samples from The COVID-19 Study

#### F.1. Study Design And Data Processing

The dataset generated from the study of Arunachalam et al. (2020) and downloaded using NCBI GEO accession number GSE152418 contained 34 subjects recruited from Atlanta, GA, USA. It included gene expression from human blood PBMC bulks samples collected from 17 healthy control subjects, one convalescent subject, and 16 subjects diagnosed with COVID-19. Healthy controls were asymptomatic adults whose samples were collected before the widespread circulation of SARS-COV-2 virus in the community. Subjects with COVID-19 diagnosis were further classified into three levels of disease severity based on the based on the adaptation of the Sixth Revised Trial Version of the Novel Coronavirus Pneumonia Diagnosis and Treatment Guidance. Moderate cases were defined as respiratory symptoms with radiological findings of pneumonia. Severe cases were defined as requiring supplemental oxygen, and ICU-hospitalized cases were those in critical conditions who needed ICU care due to organ failures.

The gene symbols were annotated to 24,259 genes using the gconvert method of the `gProfileR` package in R software. The raw counts were then converted to counts per million (CPM) using those annotated genes. Among the 16 samples from COVID-19 subjects, two (labeled S155 and S179) contained abnormally high levels of *HBB* expression (namely, *HBB* was the most highly expressed gene in those two samples; results are not shown), which could only been found in red blood cells and suggested sample contamination. Therefore, these two samples were removed from the deconvolution procedures. This also led to S155, which also provided the study with single cell samples, being excluded from the comparison in Figure 5d. The one convalescent sample was also excluded due to its unique designation.

The scRNA-seq data were generated by Arunachalam et al. (2020) using CITE-seq of >63,000 cell samples from five healthy controls and seven COVID-19 diagnosed patients. Of the seven patients, three were labeled as moderate cases, three severe cases, and one requiring ICU care. Notably, dendritic cells were enriched by the experimenters and mixed back into the samples for CITE-seq. All five healthy subjects and six of the seven (except S155, which were previously excluded) had measured bulk gene expression from independent PBMC samples as well for the analysis. To process the raw scRNA-seq data downloaded from GEO accession GSE155673, we first merged the single cell gene expression from all 12 subjects together in R package `Seurat` V4 (Hao et al., 2021). We then performed a quality control procedure for the cells, retaining those with a detected number of genes per cell between 200 and 5000 and filtering out cells with >15% mitochondrial counts. Genes with total expression less or equal to 2 across all cell samples were removed as well. We applied the global-scaling normalization method LogNormalize to normalize the feature expression measurements for each cell by the total expression, multiplied this by a scale factor of 10^6^, and log-transformed the result. We employed the UMAP dimensional reduction technique to determine the cell type notation. The R package `SingleR` (Aran et al., 2019) was used for the final determination of cell types, using gene signature data from Monaco et al. (2019) as the cell type reference. The single-cell data input now has genes as rows and estimated cell type notations with subjects as columns. In the end, 41,146 cells and 26,531 genes were retained in the gene-by-cell expression matrix of all 12 samples.

We then separated the gene-by-cell matrix into two matrices, one containing only cells from healthy controls (23,531 cells) and the other only those from COVID-19 subjects (17,615 cells). Due to the limits of memory allocations to CIBERSORTx users, 11,765 cells from healthy controls were randomly sampled for each cell type proportionally to its original relative abundance. The cell types were re-grouped by merging all cell subsets of the four major cell types of interest together. The cellular gene expression from cells not belonging to any of the four cell types were discarded for gene signature generation. The processed sub-matrices were then separately sent to CIBERSORTx software’s Create Signature Matrix module (, Newman et al., 2019) to obtain gene signature matrices for both control and COVID-19 cell samples. The signature matrix for healthy controls contains 1,368 genes for the four cell types, while that for COVID-19 samples containts 1,388 genes. After matching the common genes from the bulk expression matrices and the signature matrices returned from CIBERSORTx, there were 1,280 genes for the deconvolution of healthy control samples and 1,297 genes for the deconvolution of COVID-19 patient samples.

#### F.2. Tuning Grid And Algorithm Parameters

We adopted a tuning grid and a set of parameters for ARTdeConv to the settings in Supplementary Material Section E.3 for the analysis in Section 3.2. A random seed of 1000 was set in the R programming environment. For the deconvolution of healthy control samples, the tuning grid for *α*_1_ was set as the interval from 2^−3^ to 2^1^ with a step size of 0.2 in the power, and that for *β* was set as the interval from 2^2^ to 2^6^ also with a step size of 0.2 in the power. *α*_2_ was fixed at 10^−12^. For that of COVID-19 infected samples, the tuning grid for *α*_1_ was set as the interval from 2^−4^ to 2^0^ with a step size of 0.2 in the power, and that for *β* was also set as the interval from 2^2^ to 2^6^ also with a step size of 0.2 in the power. *α*_2_ was again fixed at 10^−12^. A 4-fold cross-validation was invoked in both deconvolution analyses to choose the optimal tuning parameters. The eventual selected tuning parameter values were within the boundaries of the grids in each case. The tolerance parameter was set as *δ* = 10^−4^.

**Figure 1:**
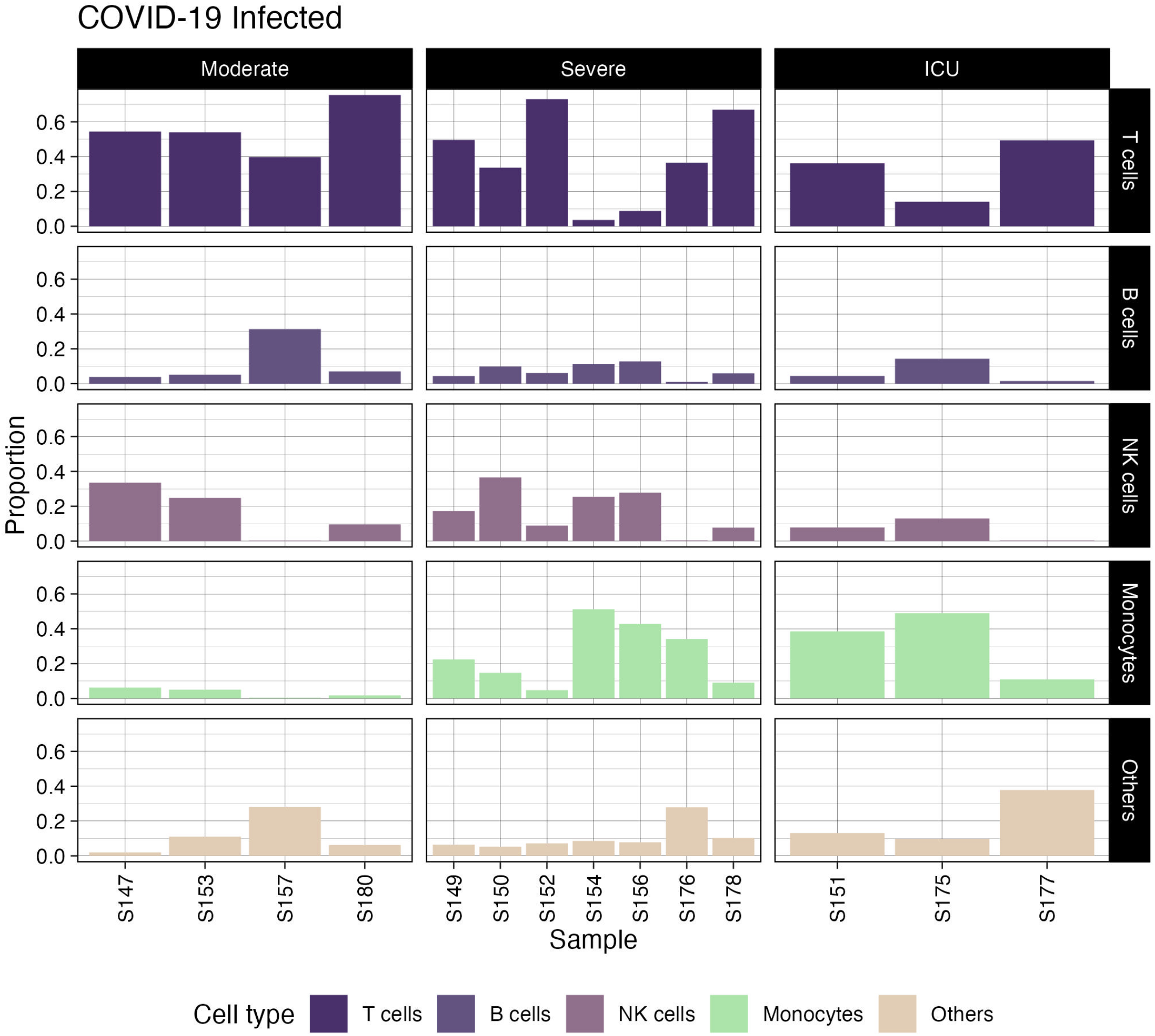
Bar graphs of the deconvolved cell type proportions of all COVID-19 infected samples, grouped by their classified disease severity

**Figure 2:**
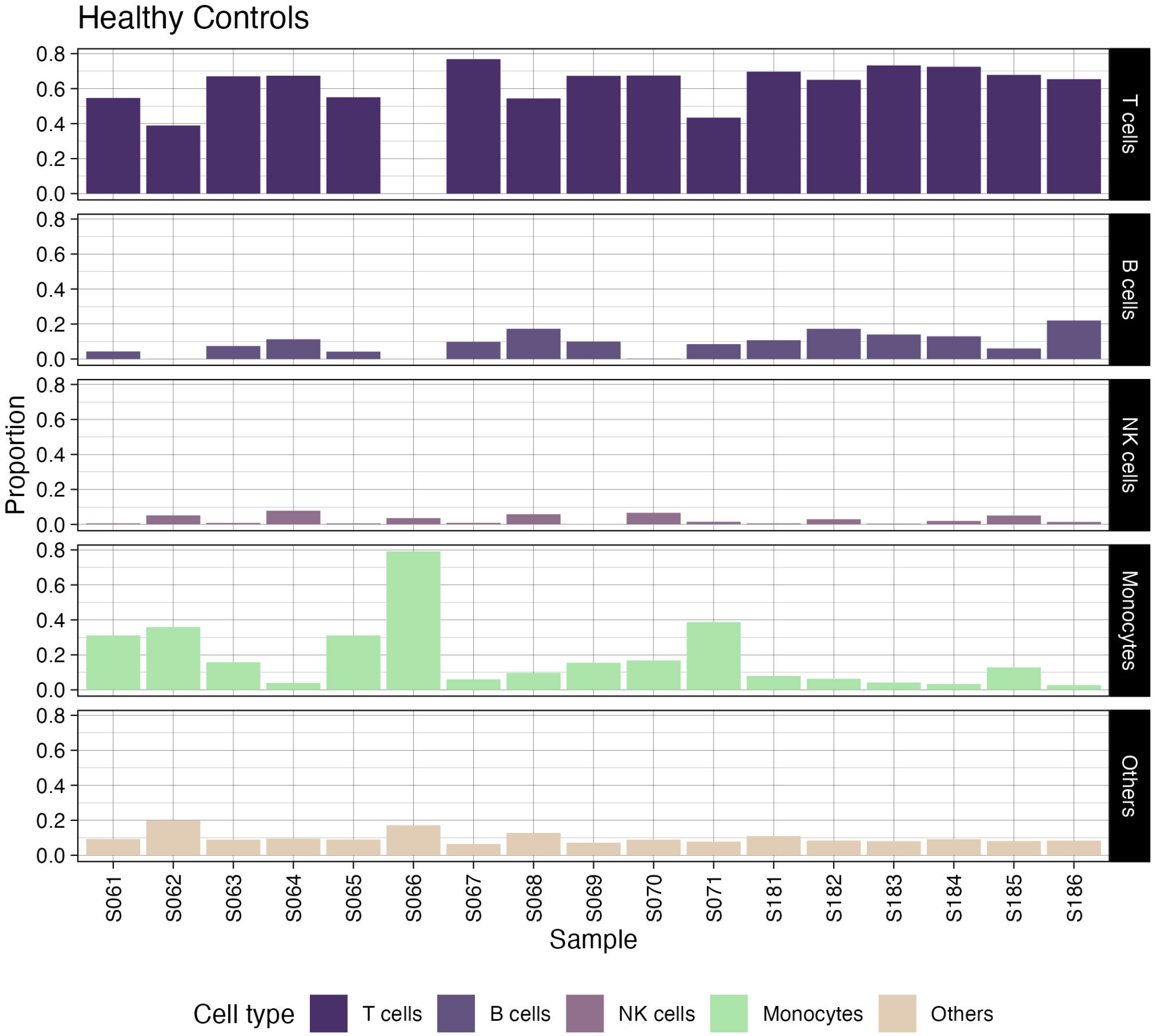
Bar graphs of the deconvolved cell type proportions of all healthy control samples.

**Figure 3:**
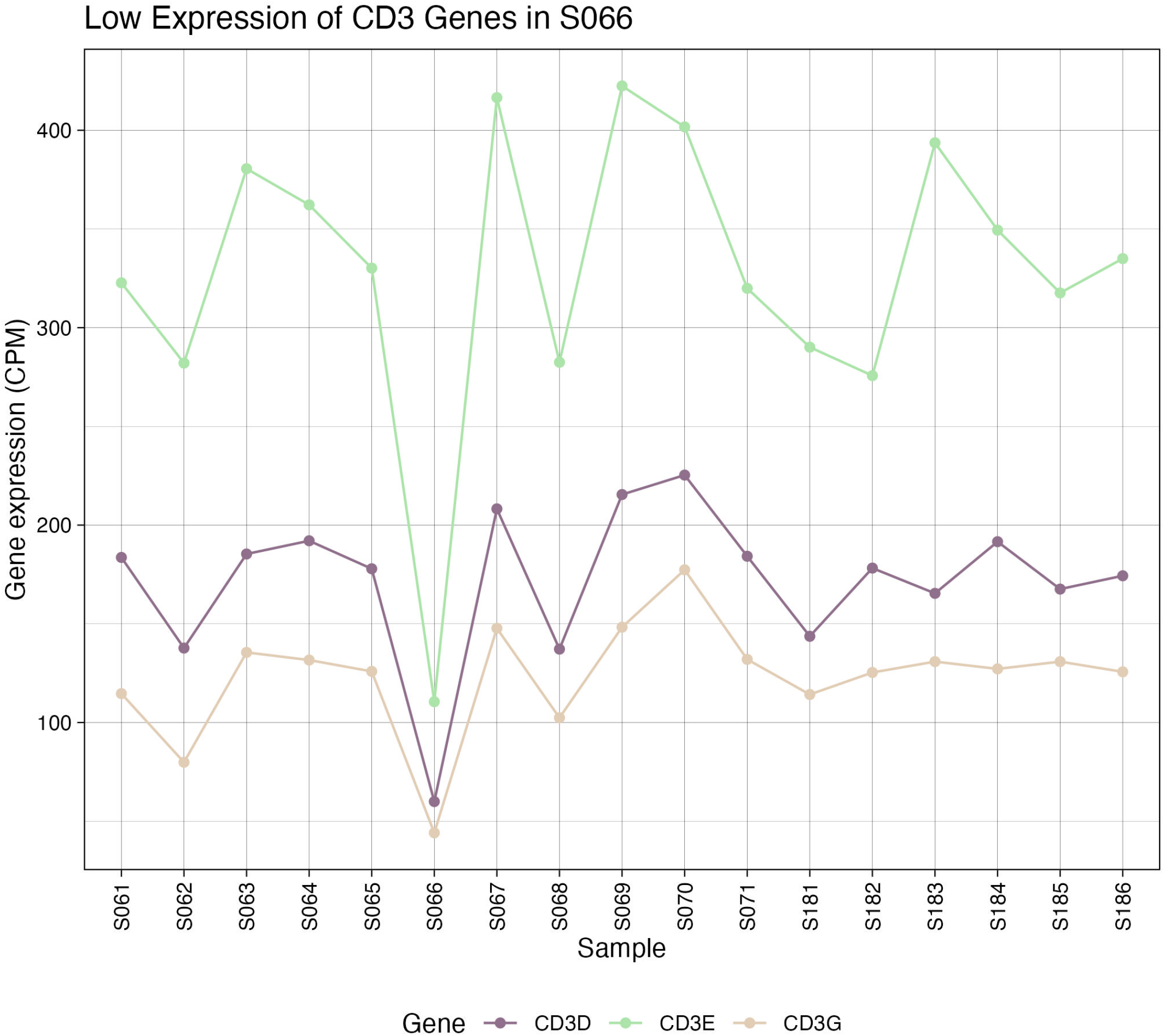
Expression of *CD3* genes, the translation of which produces T cells’ CD3 markers, among healthy samples with exceptionally low expression observed in sample S066.

**Figure 4:**
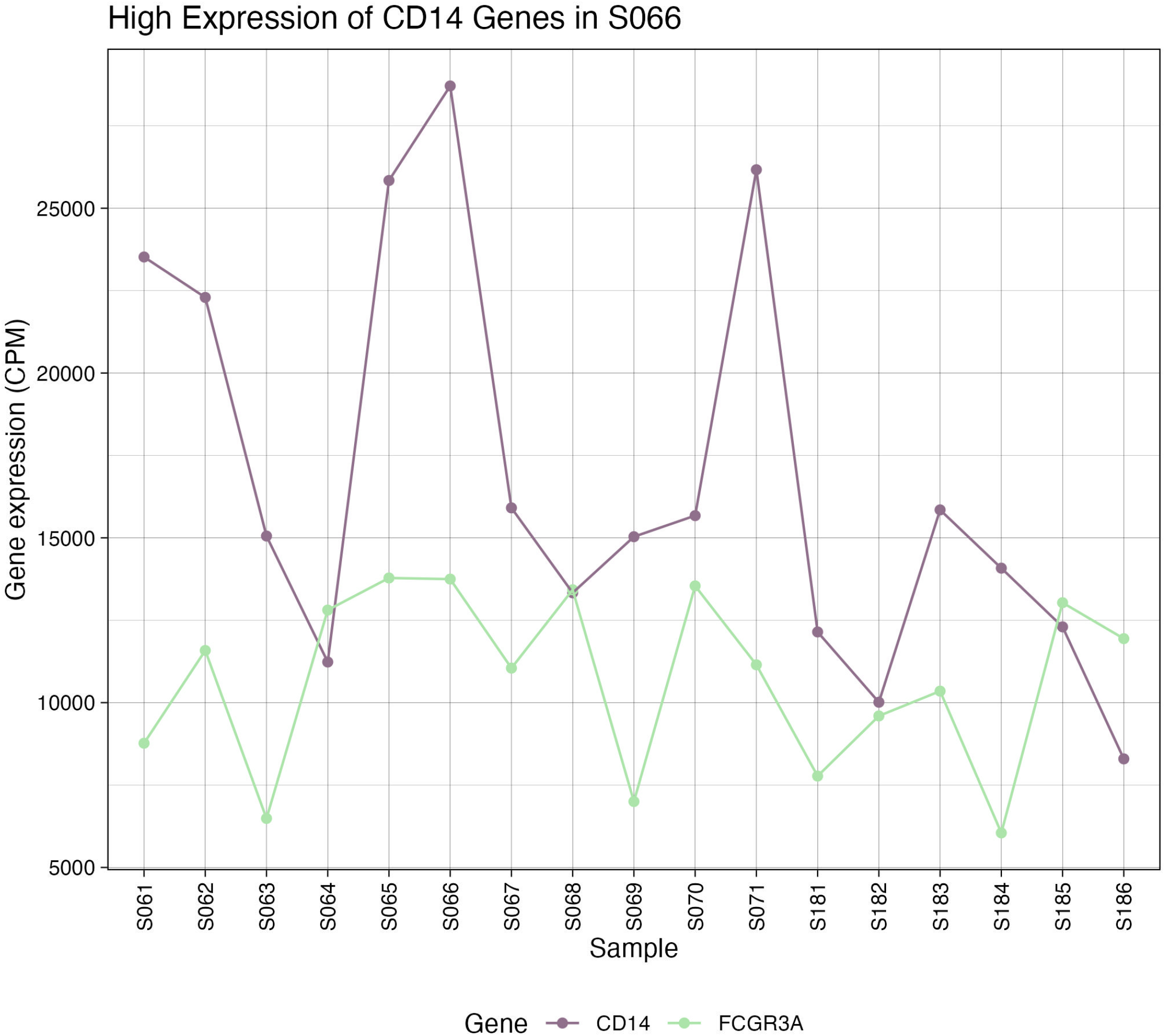
Expression of *CD14* and *FCGR3A* genes among healthy samples with high *CD14* expression observed in sample S066. *CD14* is responsible for the production of CD14 surface markers on classical/intermediate monocytes and *FCGR3A* responsible for that of CD16 surface markers on non-classical/intermediate monocytes.

